# Resilient Antarctic soil bacteria consume trace gases across wide temperature ranges

**DOI:** 10.1101/2025.09.23.677255

**Authors:** Tess F. Hutchinson, S. I. Ry Holland, David A. Clarke, Francesco Ricci, Thanavit Jirapanjawat, Pok Man Leung, Rachael Lappan, W. P. Amy Liu, Sean K. Bay, Aimee Bliss, Melodie A. McGeoch, Steven L. Chown, Chris Greening

## Abstract

Polar desert soils host diverse microbial communities despite limited nutrients and frequent temperature and light fluctuations. Adapting to these extremes, most bacteria possess high-affinity hydrogenases and carbon monoxide dehydrogenases, enabling them to use atmospheric trace gases such as hydrogen (H_2_) and carbon monoxide (CO) to generate energy and fix carbon. Despite the foundational importance of this process in polar desert ecosystems, little is known about the thermal sensitivity of trace gas oxidation or how this process will respond to climate warming. Here, we show through *in situ* and *ex situ* incubations that H_2_ consumption is an exceptionally thermally resilient process that can occur from -20 to +75°C, at rates comparable to temperate ecosystems (peaking at 8.56 nmol H_2_ h^-1^ g dry soil^-1^ at 25°C). Temperature ranges of CO (-20 to 42°C) and CH_4_ (-20 to 30°C) oxidation are also wider than expected, though the pattern of thermal sensitivity conforms with general theory. Metagenomic analyses support these data, revealing that atmospheric H_2_ and CO oxidisers are widespread, diverse, and abundant, and suggesting most Antarctic bacteria function below their temperature optima for these processes. Modelling of seasonal temperatures across ice-free Antarctica under current and future emissions scenarios indicates that H_2_ and CO oxidation can occur year-round, increasing by up to 35% or 44%, respectively, by 2100. Our results indicate constitutive aerotrophic activity contributing to Antarctic ecosystem functioning and biodiversity across spatial and temporal scales, with further studies required to understand how it interacts with photosynthesis in a changing climate.

## Introduction

The climatic conditions of Antarctica have long limited multicellular life, leading to a continent dominated by microbial primary producers [1, 2]. The small proportion (0.4%) of the continent that remains perennially ice-free is subject to temperature, light, salinity, and UV extremes, as well as a scarcity of nutrients and liquid water [3, 4]. As a result, photosynthetic primary production is highly spatially and temporally restricted throughout Antarctica [5–7]. Nevertheless, evidence is mounting that microbial communities are diverse and active even in some of the most inhospitable soils of the region [8–11]. Many of these microbes are metabolically flexible aerobes dependent on atmospheric trace gas oxidation: they can use H_2_, CO, and/or CH_4_ at rates theoretically sufficient to meet their energy needs, with H_2_ oxidation producing metabolic water as the sole end product [1, 2, 12, 13]. A proportion of these microbes can additionally fix CO_2_, thus obtaining energy and carbon required for growth from atmospheric gases, a process recently termed aerotrophy [14]. Prior to the description of this minimalistic primary production, explanations of microbial energy and carbon acquisition in Antarctica’s polar desert soils focussed on organic matter decomposition and limited autotrophic capability, falling short of accounting for the abundance of microbial biomass, especially given the paucity of photosynthetic primary producers [5, 6]. Since then, accumulating evidence from genome-resolved metagenomics, biogeochemical assays, and thermodynamic models has reinforced the significance of aerotrophy as a fundamental microbial metabolism in nutrient poor ecosystems globally [2, 12–18]. Despite this growing recognition, little is known about the thermal sensitivity or spatial distribution of this process, or its resilience to forecast climate change. Although trace gas oxidation has been reported in laboratory assays of Antarctic soils at environmentally relevant temperatures: typically 10°C, and in one instance at -20°C [1, 2, 13], the thermal limits of aerotrophy remain unknown.

Antarctic environments are characterised by substantial seasonal and daily temperature variation [19–21]. Soil temperatures in Antarctica are typically subzero but can experience extensive freeze-thaw events from austral spring to autumn, with potential for consistent positive degree days during the summer season in milder (typically, lower latitude) regions. Multiyear surface soil temperature studies have recorded maxima of 4.5°C in the McMurdo Dry Valleys, 18°C in Dronning Maud Land and Wilkes Land, and 24°C in the Antarctic Peninsula [19–22]. Soils with higher moisture, underneath insulating moss, or on north-facing slopes tend to be warmer and, particularly in coastal regions, may regularly reach 20-30°C in December and January [23, 24]. Furthermore, as global temperatures rise, Antarctica faces long-term surface warming from 0.5–3.6°C by 2100 [25] as well as heatwaves, which have caused record-breaking temperature maxima in recent years [26–28]. These events have heightened effects on soil ecosystems due to the increased heat retention of soils compared to overlying air [21].

Studies investigating the effects of long- or short-term warming on Antarctic microbial communities are scarce. They have been limited to the Antarctic Peninsula and McMurdo Dry Valleys, and predominantly focused on community composition (e.g. [29–32]), rather than function. Outside of Antarctica, numerous studies have investigated the effect of warming on soil respiration or soil organic carbon degradation, showing that the microbial population is an important driver of soil temperature sensitivity [33, 34]. However, the growth and function of each microbe will have unique thermal responses [35], making it difficult to predict the emergent properties of a soil ecosystem under warming. These ecosystem-scale investigations have been complemented by culture-based studies, which have often demonstrated Antarctic isolates capable of, or even optimally growing at, mesophilic temperatures (15-30°C, e.g. [36–38]). The few functional studies in Antarctic soils have shown increases in soil respiration [39, 40] and degrading enzyme activities [41, 42] with temperature, yet trace gas oxidation remains unexplored.

Given the key role of aerotrophy in sustaining the biodiversity and productivity of Antarctica, it is critical to understand the thermal responses of this process. To address this, we measured trace gas oxidation both *in situ* at natural summer temperatures and *ex situ* across a wide range of temperatures (-20 to 80°C) in three major Antarctic regions, in conjunction with genome-resolved metagenomic profiling. We then tested whether a purported general theory for biological temperature dependence [43] extends to this data and used continent-wide modelling to predict consumption rates for H_2_ and CO across Antarctic Conservation Biogeographic Regions (ACBRs) under current and future climates. This is the first study investigating the effect of warming on soil microbial ecosystems in Dronning Maud Land or Wilkes Land, which contain significant ice-free oases. Our findings suggest that thermally resilient microbes mediate trace gas oxidation year-round in Antarctic soils and indicate that this process is likely to increase with forecast warming scenarios.

## Materials and Methods

### Sampling locations

Samples were collected in austral summers from several ice-free regions of continental Antarctica. Soil samples from Dronning Maud Land were collected from the Schirmacher Oasis, Henriksen Nunataks, and Holtedahl Mountains in January 2024, and from the Petermann Ranges and Humboldt Mountains in December 2024 (Fig. 1A). Schirmacher Oasis is a 34 km^2^ region on the Princess Astrid Coast that lies between the ice shelf and the East Antarctic Ice Sheet; whereas the other sites lie further inland in the Antarctic slopes climatic zone [44]. The nunataks and mountain ranges are characterised by rocky outcrops covered in dry sediments, often with a layer of small rocks and pebbles on the surface. See Table S1 for full details of all sampling locations.

**Figure 1.**
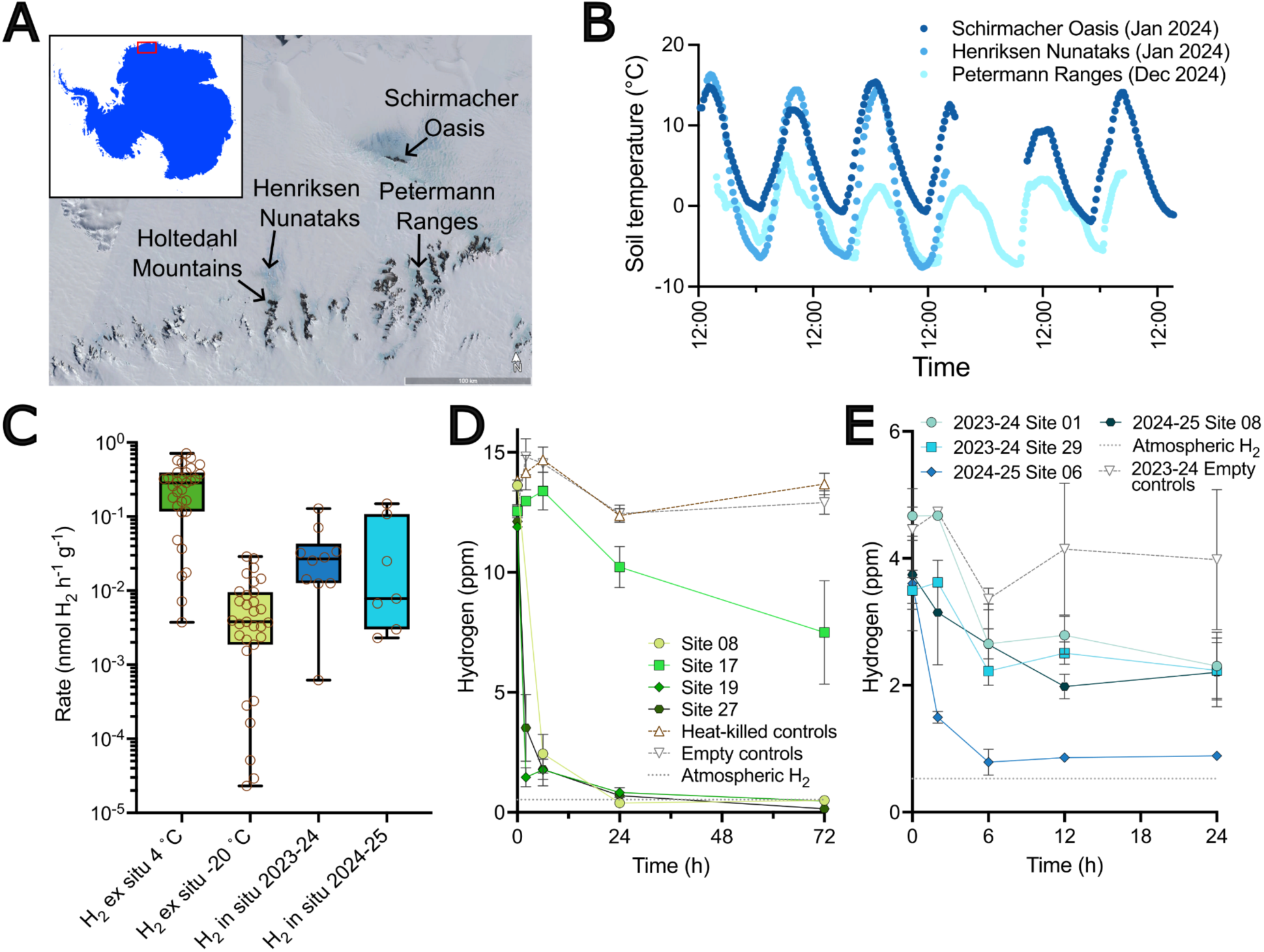
Hydrogen oxidation is a widespread trait across Dronning Maud Land, Antarctica, occurring in *in situ* and *ex situ* soil microcosms. (**A**) Map of broad sampling locations across Dronning Maud Land, Antarctica. Inset shows entire continent, with main map area boxed in red. Satellite imagery is from Google Earth. (**B**) Daily soil temperature variation from the *in situ* temperature probe. The x axis represents time (24 h format). Readings were taken over different time periods but are shown overlaid. (**C**) Boxplot of H_2_ consumption rates in Dronning Maud Land soil microcosms incubated *ex situ* at 4 and -20 °C (both *n* = 31) and *in situ* in the 2023-2024 (*n* = 10) and 2024-2025 (*n* = 7) seasons. Rates are presented on a log scale. Boxplots show means and full range of data; sample values are shown as brown circles. (**D**) H_2_ consumption for 4 representative samples in *ex situ* microcosms incubated at 4 °C (*n* = 3 ± SD). (**E**) H_2_ consumption for 4 representative samples in *in situ* microcosms (*n* = 3 ± SD). Trace gas oxidation rate calculations for panels (CDE) are in Table S2.

Robinson Ridge sits onshore from the Windmill Islands in Wilkes Land (Fig. 2A). The site is characterised by rocky outcrops covered in cobble-sized gravel interspersed with small boulders. Lichens were prevalent throughout, although restricted to larger gravel rocks, which were removed before sampling. Soil samples were collected in February 2022. The Bunger Hills is a ∼450 km^2^ region that lies to the east of Robinson Ridge, also within Wilkes Land (Fig. 2A). It is one of the largest ice-free regions in Antarctica outside of the McMurdo Dry Valleys and features a series of coastal hills interspersed with numerous small, predominantly saline lakes and ponds [45], as well as several large offshore islands to the north. Bunger Hills soil samples were collected between December 2023 and January 2024 (Table S1).

**Figure 2.**
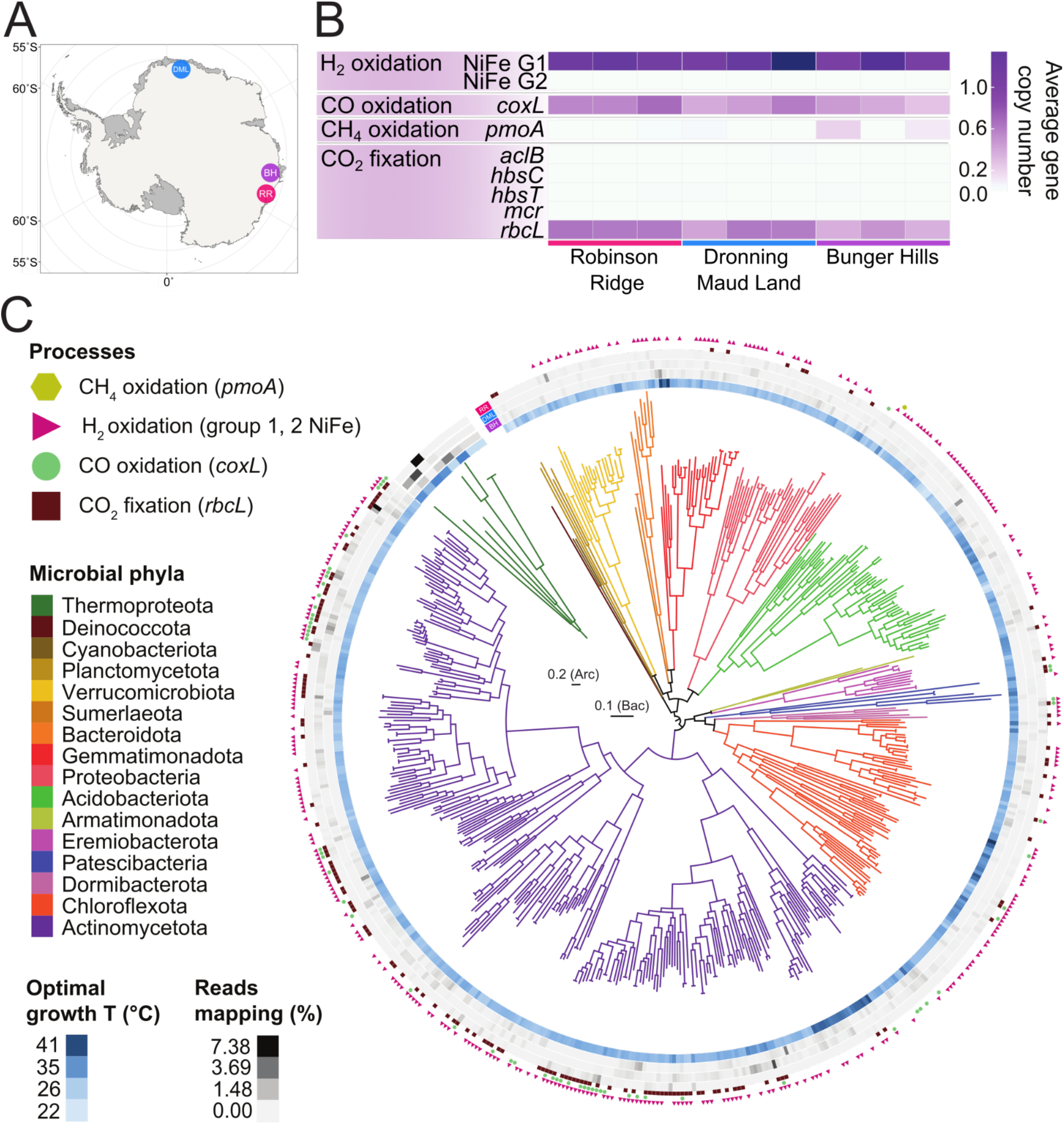
Trace gas oxidation genes are widespread across Antarctic metagenomes and multiple phyla. (**A**) Map of sampling locations across Antarctica. DML, Dronning Maud Land; BH, Bunger Hills; RR, Robinson Ridge. (**B**) Average gene copy numbers per metagenome of metabolic marker genes mediating carbon fixation and trace gas oxidation in the Antarctic soil microbial communities. (**C**) Archaeal and bacterial maximum likelihood phylogenetic trees of the recovered 554 MAGs, including their optimal growth temperature, percentage of location-based metagenome reads that map to each MAG, and the presence/absence of key genes for atmospheric trace gas oxidation and carbon fixation through the Calvin Benson Bassham cycle.

### Sampling methods

In all locations, soil samples (100 - 500 g) were collected from the top 0-10 cm of the soil surface after clearing any larger pebbles or rocks, using a sterilised stainless steel trowel and stored in sterile Whirl-Pak sample bags. The trowel and nitrile gloves worn by sampling personnel were sterilised with 70% isopropanol wipes immediately before sampling. At Dronning Maud Land sites, triplicate samples were collected from within a 1 m^2^ plot and analysed as biological replicates throughout the experimental work. Samples were collected from a total of 48 plots (Robinson Ridge *n* = 3, Bunger Hills *n* = 3, Dronning Maud Land *n* = 31 in 2023-24 and *n* = 11 in 2024-25). Soil samples were kept below 0°C for the duration of the field campaigns, before being transported to Monash University, Australia at -18°C, then stored at the same temperature until further processing and analysis.

At three separate field camps in Dronning Maud Land, soil temperature was measured with a ZL6 data logger equipped with a TEROS 12 soil moisture and temperature sensor (METER Group, Inc., Washington, USA). The temperature sensor has an accuracy of ± 1.0°C below 0°C or ± 0.5°C above 0°C. At each location, the sensor was buried at 5-10 cm depth (Fig. S1ABC), remained in the ground for 2-7 days and the logger recorded data every 10 or 30 minutes.

### Community DNA extraction and sequencing

DNA was extracted from technical-duplicate 0.5 g soil samples for three sites each from Dronning Maud Land, Robinson Ridge, and Bunger Hills (total *n* = 9) using the FastDNA Spin Kit for Soil (MP Biomedical, California, USA) with a modified protocol for low biomass, in which the technical-duplicate soil samples were combined during the binding matrix step, then washed and eluted as one. DNA was quantified via a Qubit High Sensitivity Assay for dsDNA. A negative control in the form of a DNA extraction blank without any sample was included in sequencing and analysis. The extracted DNA was submitted to the Monash Genomics & Bioinformatics Platform (Melbourne, Australia) for library preparation and sequencing, which was carried out on a single lane of a NovaSeqX Plus (Illumina) using XLEAP-SBS chemistry (2 × 150 bp). The negative control had 316 read pairs, whereas the nine soil samples had an average of over 50 million read pairs (range 39,404,210 - 63,902,121).

### Metagenomic analysis

Quality control, assembly, and binning of the reads were performed using the Metaphor pipeline [46]. Raw reads from the nine metagenomic libraries and control sample underwent quality filtering with fastp [47], which included trimming of primers and adapters and removal of artefacts and low-quality reads with parameters *length_required = 50*, *cut_mean_quality = 30*, and *--detect_adapter_for_pe*. The resulting high-quality reads were coassembled using MEGAHIT v1.2.9 [48] with default settings. Coassembly was selected due to the similar nature of the samples, aiming to improve recovery of low-abundance microbial taxa and reduce contig redundancy. Contigs shorter than 1,000 bp were discarded. Taxonomic profiling of the metagenomes was performed with phyloFlash [49].

To generate metagenome-assembled genomes (MAGs), binning was conducted using Vamb v4.1.3 [50], MetaBAT v2.12.1 [51], SemiBin v2.2.0 [52], and CONCOCT v1.1.0 [53]. The resulting bins were refined with the bin refinement module of MetaWRAP [54] and dereplicated using dRep v3.4.2 [55] at 99% ANI, incorporating quality assessment from CheckM2 v1.1.0 [56]. CheckM2 v1.1.0 was also used to estimate completeness and contamination of all bins. Following dereplication, we obtained 554 MAGs of medium (completeness >50%, contamination <10%) to high quality (completeness >90%, contamination <5%), following MIMAG standards [57]. MAG taxonomy was assigned using GTDB-Tk v2.3.2 [58] based on Genome Taxonomy Database Release 08-RS214 [59]. Relative abundances of the MAGs across samples were calculated using CoverM v0.6.1 [60] in genome mode. Maximum-likelihood phylogenetic trees for archaeal and bacterial MAGs were constructed using the identify and align commands in GTDB-Tk [58]. Genome trees were inferred with IQ-TREE v2.3.6 [61, 62] using 1,000 ultrafast bootstrap [63] replicates, and the Q.insect+F+R3 and LG+F+R10 substitution models for archaeal and bacterial datasets, respectively. Phylogenetic trees were visualised with iTOL v6 [64] and finalised in Illustrator v24.0.2 (Adobe Inc., California, USA).

### Metabolic annotations

Metabolic marker genes spanning pathways mediating energy conservation, carbon fixation, trace gas metabolism, the sulfur and nitrogen cycles, arsenic and iron cycling, formate oxidation, phototrophy, and aerobic respiration were identified from the 554 MAGs. Open reading frames were predicted using Prodigal v2.6.3 [65], followed by contig annotation through homology-based searches using DIAMOND blastp [66] against a custom database [67] comprising 51 curated metabolic marker gene sets (described below). Temperature optima for MAGs were predicted using Tome [68].

Metabolic analysis of the entire microbial community was also carried out with the short-read metagenomic data. Using the BBDuk function within BBTools v36.92 (https://sourceforge.net/projects/bbmap/) paired-end reads from the nine samples and the negative control were stripped of adapter and barcode sequences, and contaminating PhiX reads as well as low-quality sequences (minimum quality score of 20) were filtered out. High-quality forward reads of at least 130 bp length were subsequently screened for the 51 metabolic marker genes using the DIAMOND blastx algorithm [69]. Reads were aligned against the custom reference database [67], then alignments were filtered using a minimum query coverage of 80% and identity thresholds of 80% for *psaA*, 75% for *hbsT* (contigs: 65%), 70% for *atpA* (contigs: 60%)*, psbA* (contigs: 60%)*, isoA, ygfK*, and *aro*, 60% for *amoA, mmoA, coxL*, [FeFe]-hydrogenase, *nxrA, rbcL*, and *nuoF*, and 50% for all other genes (contigs: *rho* 30%; *rdhA*: 45%; *cyc2*: 35%). To estimate the proportion of community members encoding each gene, read counts were normalised to reads per kilobase per million (RPKM) and further scaled by the average RPKM of 14 universal single-copy ribosomal marker genes.

### Trace gas oxidation assays

Three different sets of trace gas oxidation assays were performed: (1) *in situ* microcosms incubated at field camps in Dronning Maud Land over the 2023-24 and 2024-25 field seasons; (2) *ex situ* microcosms with all soils collected from Dronning Maud Land in the 2023-24 field season at representative summer (4°C) and winter (-20°C) temperature; and (3) *ex situ* microcosms with three representative soils each from Dronning Maud Land, Robinson Ridge, and Bunger Hills at 14 different temperatures. For all incubations, soil was aliquoted into 120 ml glass serum vials (Wheaton Science Products, New Jersey, USA), sealed with NaOH-rinsed butyl rubber stoppers and aluminium crimp caps before an equal mixture of H_2_, CO, and CH_4_ (procured as a premixed gas in N_2_; BOC) was added to an ambient air headspace to achieve a final headspace concentration of 10 ppm.

For *in situ* Dronning Maud Land incubations, 5 g of a single replicate soil from select sites was added to vials in triplicate (i.e. the biological triplicates were not used here). Serum vials were prepared, incubated and sampled outdoors in ambient Antarctic conditions (surface air temperature ∼-18 to 5°C) and 10 ppm gas mix was added to empty, sealed vials in duplicate or triplicate for negative controls. Serum vials were secured in a custom rack staked into either soil or snow, depending on the location of the field camp (Fig. S1DE).

For laboratory-based *ex situ* incubations at 4 and -20°C as representative summer and winter temperatures, respectively, biological triplicate soil samples from all 31 Dronning Maud Land sites sampled in the 2023-24 field season were prepared under sterile conditions. Serum vials with soil inside were left to equilibrate at the incubation temperature for at least two hours before they were sealed and injected with the same gas mix described above. Heat-killed soil samples (*n* = 3, randomly chosen sites) and empty serum vials (*n* = 3) were both used as negative controls.

For laboratory-based, *ex situ* temperature range incubations, three soil samples from each of Robinson Ridge, Dronning Maud Land, and the Bunger Hills (single replicate of each, nine soils total) were prepared as above and incubated at 14 different temperatures (-20, -8, 4, 10, 17, 25, 30, 37, 42, 50, 60, 70, 75 and 80°C). For Dronning Maud Land and Bunger Hills, the soil samples were chosen from environments to ensure maximal possible geographic spread based on the samples taken. One pooled heat-killed soil sample per region was used as a control at each temperature. Soil (1-10 g) was prepared and left to equilibrate for two hours at the incubation temperature, before the vials were sealed and mixed gas was added to a final concentration of 10 ppm, as described above.

For all incubations, at each time point, 2 ml gas samples were withdrawn via a syringe fitted with a stopcock and stored in 3 ml glass exetainers sealed with silicone and flushed with N_2_ as described previously [70]. Gas sample exetainers were analysed on a VICI gas chromatograph with a pulse discharge helium ionisation detector as previously described [71] and quantified by comparison to a three-point standard curve (0, 10, 100 ppm each of H_2_, CO, CH_4_). Previously reported microcosms containing Mackay Glacier soil samples [2] were also resampled for a 4-year timepoint by manual injection into the same instrument and similarly compared to a three-point standard curve (0, 0.5, 10 ppm CH_4_). Trace gas oxidation rates were determined by fitting exponential growth curves to the depleting gas concentrations over time, normalised to controls.

Trace gas consumption rates per gram of dry soil were determined by first quantifying gravimetric soil moisture content: quantified by calculating the weight difference (to 4 decimal places) of 5 g soil after drying at 70°C for one week, followed by 105°C for seven hours. Samples were placed in a desiccator to return to room temperature before weighing. Cell specific gas consumption rates were also calculated for the nine soil samples used in the *ex situ* temperature range incubations, by dividing bulk oxidation rates by the number of cells capable of each trace gas oxidation in the soil as determined by 16S rRNA gene qPCR (following a previously described method [2]) and metabolic marker copy number. Normalisation to 16S rRNA and metabolic marker genes was performed as in [2]. The average metabolic marker gene copies per organism was derived from short read analysis of the nine metagenomes.

### Oxidation rate models

To model the thermal sensitivity of atmospheric trace gas oxidation, a universal theory of temperature dependence in biology [43] was applied to data from the *ex situ* incubations of nine soils at 14 different temperatures. This had a secondary aim of testing whether this general theory also applied to the unique polar desert ecosystems of Antarctica, where it h ad not previously been tested. Model equations from [43] were applied in R v4.4.0 to determine fit, optimum temperature (*T*_opt_), and thermodynamic parameters (ΔC, ΔH).

Following this, to predict the temporal and spatial distribution of the process under current and future climates, zero-inflated generalised linear models (GLM) with a gamma error distribution were fitted to bulk H_2_, CO and CH_4_ oxidation rates. A zero-inflated approach was used to mitigate the relatively large amounts of zeroes in the data, which are incompatible with a gamma distribution. This was preferred over the decision to add small amounts of noise to each zero value given the proportion of zeros (24% H_2_, 50% CO, 81% CH_4_) and the already low non-zero values. Models used a second-order polynomial for temperature and evaluated both conditional (i.e. using only non-zero rate values) and zero-inflated components. Models were fitted using the glmmTMB function from the glmmTMB R package [72] whereby the family was specified as a zero-inflated gamma with a log link function. For each gas, temperature (°C) was included as a second-degree polynomial to account for the non-linear association with oxidation rate. This was the case for both the gamma and zero-inflated components of the model. Model fit and diagnostics were assessed using the simulateResiduals function from the DHARMa R package [73], with *n* = 1000 simulations (Fig. S2). To assess prediction accuracy of the models, and account for sample size, we performed *n*-fold, or leave-one-out, cross validation using the cv R package [74]. Mean squared error was used as the criterion for estimating prediction accuracy.

The GLMs were then used to estimate spatial variation in oxidation rates across the continent, using high resolution climate data for the land surface. Temperature at surface data for use in predicting rates of trace gas oxidation across ice-free Antarctica were obtained from CHELSA v2.1 as raster layers at a ∼ 1 km resolution [75, 76]. All climatologies from CHELSA v2.1 were used. This included four time periods (1981-2010, 2011-2040, 2041-2070, 2071-2100) where all but 1981-2010 (“baseline” climate) are associated with outputs from five global circulation models (GFDL-ESM4, IPSL-CM6A-LR, MPI-ESM1-2-HR, MRI-ESM2-0, UKESM1-0-LL), and three emissions scenarios (ssp126, ssp370, ssp585) per model. Surface air temperatures were used in the absence of soil temperatures, which will likely cause some underestimation of rates, given soils are typically warmer. The model also assumes the three sampled regions are generally representative of all ice-free areas; the low prediction error rates (see results) support this, though further data are needed for full validation. All temperature data were cropped so as to only include data south of 50°S. This was then re-projected from WGS84 to the ESRI:102020 coordinate reference system, which is an equal-area polar azimuthal projection. For each emission scenario and time period, we used the mean of the five global circulation models when making the spatial predictions. Finally, the Antarctic Conservation Biogeographic Regions (ACBR) [77, 78] spatial layer was used to mask the final predictions, given that we were only interested in oxidation rates within the ice-free areas of Antarctica.

## Results

### Soil microbial trace gas oxidation occurs *in situ* and *ex situ* in Dronning Maud Land

We conducted a comprehensive sampling campaign of Dronning Maud Land, collecting 31 surface soil samples from the coastal Schirmacher Oasis, inland nunataks, and mountain ranges (Fig. 1A, Table S1). Through *ex situ* incubations at 4°C, chosen as a representative summer temperature, all 31 soils consumed H_2_ (28 of these to sub-atmospheric levels; below 0.53 ppm) within the eight-week incubation (Fig. 1CD). Of these 4°C incubations, 25 also consumed CO and 12 consumed CH_4_ (three to sub-atmospheric levels, below 1.9 ppm; Fig. S3). Bulk rates of H_2_ oxidation varied by several orders of magnitude, from 3.6 × 10^−3^ to 7.1 × 10^−1^ nmol H_2_ h^-1^ g dry soil^-1^ (Fig. 1C). At the representative winter temperature (-20°C), 21 of the 31 soil samples still consumed H_2_ to sub-atmospheric levels, with bulk rates again spanning two orders of magnitude, from 2.3 × 10^-5^ to 2.9 × 10^-2^ nmol H_2_ h^-1^ g dry soil^-1^ (Fig. 1C). Two soil samples also consumed CO, and five consumed CH_4_ at this temperature, which represents the first report of the latter metabolic process occurring at subzero temperatures (Fig. S3). Microbial communities where this activity occurred may therefore be capable of growth, or at least energy-harvesting for cellular maintenance, year-round. Overall, these results confirm that trace gas oxidation is a general process that is preserved across the landscapes of Dronning Maud Land, despite their harsh and varied physicochemical conditions. Moreover, the oxidation of H_2_, CO and, to a lesser extent, CH_4_ is preserved at temperatures chosen as representative of summer (4°C) and winter (-20°C) conditions for the region.

To further validate that trace gas oxidation activity persists under environmental conditions in Antarctica and is not an artifact of sample handling or storage, freshly collected soils from Dronning Maud Land were analysed for their ability to oxidise trace atmospheric gases *in situ* during the 2023-24 and 2024-25 field seasons. H_2_ oxidation was observed in 16 of the 17 topsoil samples at an average rate of 3.83 × 10^-1^ nmol H_2_ h^-1^ g dry soil^-1^ (95% CI: 8.8 × 10^-2^ to 6.8 × 10^-1^, *n* = 10) in 2023-24 and 3.0 × 10^-1^ nmol H_2_ h^-1^ g dry soil^-1^ (95% CI: 7.0 × 10^-2^ to 5.3 × 10^-1^, *n* = 7) in 2024-25 (Fig. 1CE). Although measured H_2_ did not reach below sub atmospheric levels in the *in situ* incubations (Fig. 1E), this was likely due to technical constraints and abiotic gas emission from stopper materials due to the storage of gases at low concentrations in the 3 ml exetainers over a period 4-6 weeks [70]; exetainer samples are usually analysed within 48 hours during *ex situ* incubations, but samples from the *in situ* microcosms could not be analysed until the field team returned to Australia. Oxidation of CO and CH_4_ during the *in situ* incubations was not conclusively observed (Fig S4). This may reflect abiotic production of CO from the gas storage exetainers over this extended period of time [70] and the slow or absent rates of CH_4_ oxidation at low temperatures, also observed in the *ex situ* incubations (Table S2).

Nonetheless, the Dronning Maud Land soil microbial communities oxidised H_2_ at rates of comparable magnitude in the *in situ* and *ex situ* incubations, accounting for the temperature differences (Fig. 1CDE). Soil temperature loggers placed at 10 cm soil depth in both the 2023-24 and 2024-25 seasons confirmed that soils in the region varied in temperature from -1.9 to 15°C at Schirmacher Oasis (mean 5.0 °C, January 2024), -7.6 to 16°C at Henriksen Nunataks (mean 2.5 °C, January 2024), and -7.2 to 6.3°C in the Petermann Ranges (mean -1.1°C, December 2024) (Fig. 1B). These data add to previous soil temperature recordings from Dronning Maud Land, as none were available for the specific inland nunatak and mountain regions visited in these expeditions.

### High capacity for trace gas oxidation across geographically disparate Antarctic communities

To profile the capacity and activities for trace gas oxidation across a wider geographical range, we next investigated polar desert soils from three disparate locations in East Antarctica: Dronning Maud Land, Robinson Ridge, and Bunger Hills (*n* = 3 in each location, Fig. 2A, Table S1). We used genome-resolved metagenomics to investigate microorganisms with the capacity for atmospheric trace gas oxidation across these distant, disconnected, and understudied ice-free regions. Analysis of metagenomic short reads confirmed that the capacity for aerotrophy was widespread across all nine samples (Fig. 2B, Table S3). Assembly and binning resulted in the construction of 554 medium-to high-quality strain level metagenome-assembled genomes (MAGs), spanning 15 bacterial and 1 archaeal phylum (Fig. 2C, Table S4). Community composition was similar across the soils, with *Actinomycetota* dominant (283 MAGs, 29.5 ± 11.5% metagenomic short reads), and *Chloroflexota* (71 MAGs, 3.5 ± 1.0% reads) and *Acidobacteriota* (60 MAGs, 3.3 ± 0.6% reads) also prevalent (Fig. 2C, Fig. S5, Table S5). Maximum likelihood phylogenetic trees of the 554 MAGs show the diversity within each phylum, especially within dominant *Actinomycetota* (Fig. 2C). Read mapping showed that although some taxa were distributed across all three locations (78 MAGs), most were range-restricted to just one of Dronning Maud Land, Bunger Hills, or Robinson Ridge (Table S5). Ubiquitous taxa included some common soil lineages (e.g. *Mycobacterium*, *Nocardiodes*, *Streptomyces*, *Solirubrobacter*), as well as genera that include psychrophilic species isolated from tundra and permafrost (*Abditibacterium*, *Spirochaeta* [79, 80]) or capable of growth at cold temperatures (*Crossiella* [81]). Genome-based *in silico* prediction of the optimal growth temperature for each MAG revealed values between 22-43°C (median 31°C, Fig. 2C, Table S4). This indicates that a large proportion of the microbial community could be functioning in conditions far below their optima for growth and will persist even under climate warming scenarios.

Analysis of metabolic genes in the metagenomes and MAGs confirmed that the capacity for trace gas oxidation was widespread in the sampled soils and across phyla. The genes for atmospheric H_2_ oxidation (group 1h and 1l [NiFe]-hydrogenase large subunits) were encoded by over half of the MAGs (*n* = 234 and *n* = 119, respectively, inclusive of 31 MAGs encoding both), spanning 11 of the 15 phyla (Fig. 2C). These genes were predicted to be encoded by most bacteria based on metagenomic short reads (>1 average gene copy number per organism at most sites) (Fig. 2B, Table S3). There was also considerable capacity for CO oxidation (CO dehydrogenase catalytic subunit *coxL*; 60 MAGs from 5 phyla, 0.49 average copies) and chemosynthetic CO_2_ fixation (type IE RuBisCO catalytic subunit *rbcL*; 120 MAGs from 8 phyla, 0.52 average copies) (Fig. 2C). As 44 of the 60 MAGs containing *coxL* genes also encode group 1h or 1l [NiFe]-hydrogenases, H_2_ oxidation is likely the preferred energy source in Antarctic soils, perhaps due to the cellular hydration potential the process provides in arid environments [2, 13]. As only 97 of the 339 MAGs encoding aerotrophy genes (a high-affinity group 1h or 1l hydrogenase, CO dehydrogenase, or methane monooxygenase) also encoded a carbon-fixing type IE RuBisCO gene (Table S4), the remaining MAGs may instead rely on heterotrophic carbon sources, such as scavenging necromass, or use atmospheric trace gases primarily to support maintenance and persistence rather than growth. In contrast to the high capacity for H_2_ and CO oxidation, only one MAG (taxonomically assigned to atmospheric methanotroph genus *Methylocella*) harboured the *pmoA* gene for atmospheric methane oxidation (Fig. 2C, Table S4). Correspondingly, this gene was only abundant in two Bunger Hills samples (Fig. 2B). Results from the Dronning Maud Land *ex situ* incubations suggest that CH_4_ oxidation may be restricted to coastal regions with milder weather, as this trait was observed in some soils from the Schirmacher Oasis, but not in any soil samples from the colder, high altitude inland mountain ranges or nunataks (Table S1,2).

### Antarctic soils sustain trace gas oxidation across a wide range of temperatures

To assess the thermal sensitivity of atmospheric trace gas oxidation in Antarctic soils, the surface soils used for metagenomic analyses from Robinson Ridge (*n* = 3), Dronning Maud Land (*n* = 3) and Bunger Hills (*n* = 3) were incubated in *ex situ* microcosms at 14 different temperatures. These ranged from -20 to 80°C and encompassed temperatures relevant to Antarctic soil conditions (-20, -8, 4, 10, 17, 25°C) and the predicted optimal growth temperatures for the MAGs (30, 37, 42, 50°C). H_2_ oxidation was ubiquitous, with most soils displaying a wide thermal tolerance from -20 to 50°C (Fig. 3A, Table S6). The highest rates of oxidation were consistently observed between 17 and 37°C (Fig. 3AB), where H_2_ (10 ppm) was rapidly consumed to sub-atmospheric levels (<0.53 ppm), often within 1 hour (Fig. S6). To explore the upper thermal limit of H_2_ oxidation, soils were also incubated at temperatures of 60 to 80°C. All soils could oxidise H_2_ at 50°C, but only some Dronning Maud Land and Bunger Hills soils could maintain this at higher temperatures of 60 and 70°C. Bunger Hills soils were even capable of H_2_ oxidation at a maximum of 75°C for ∼24 hours, after which activity ceased, presumably due to cell death (Fig. S7). Despite the high thermal tolerance of the hydrogenotrophic community, predicted optimal growth temperatures of the MAGs encoding group 1h and 1l [NiFe] hydrogenases peaked at 43.9°C (JACCZL01; a member of the phylum *Chloroflexota*, Table S4).

**Figure 3.**
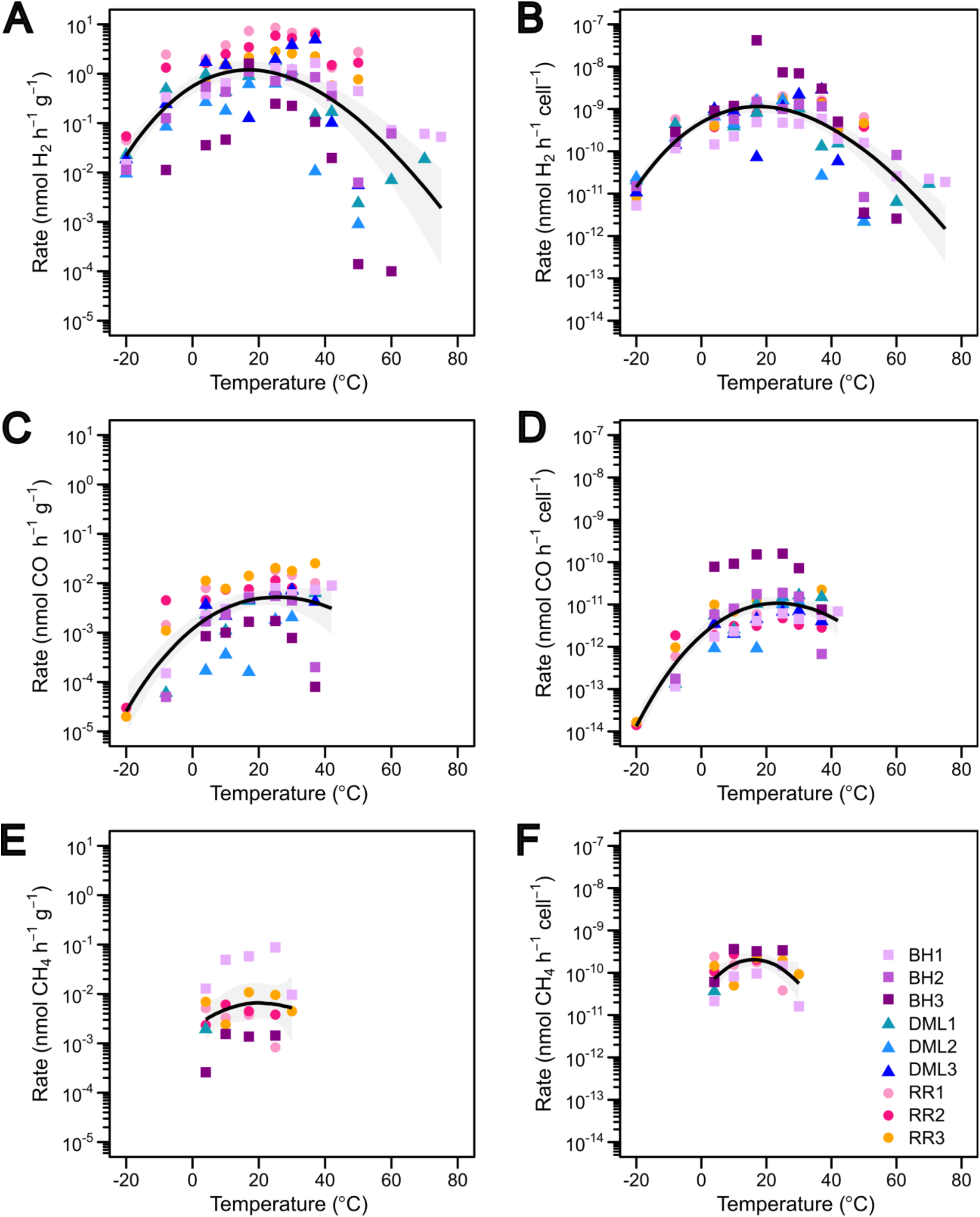
Antarctic soils oxidise trace atmospheric gases across a wide range of temperatures. Rates of H_2_, CO and CH_4_ oxidation are presented as both bulk oxidation rates (nmol h^-1^ g^-1^, **A,C,E**) and cell-specific oxidation rates (nmol h^-1^ cell^-1^, **B,D,F**). The General Temperature Dependence (GTD) model was used to generate a fit line, with a 95% confidence interval shown in grey. RR, Robinson Ridge; DML, Dronning Maud Land; BH, Bunger Hills; MG, Mackay Glacier.

Both CO and CH_4_ oxidation displayed a narrower thermal range and slower rates than for H_2_ oxidation. CO oxidation occurred in all soils at temperatures of -20 to 42°C, with bulk soil and cell-specific rates consistently ∼2 orders of magnitude lower than for H_2_ (Fig. 3CD, Table S6). Beyond 42°C, accurate quantification of CO was obstructed by abiotic production of the gas due to thermal degradation of butyl rubber stoppers at increased temperatures [70, 82]. CH_4_ oxidation was only observed in some soils from 4 to 30°C (Fig. 3E, Table S6), consistent with the sporadic distribution and relatively low abundance and diversity of methanotrophs reported via metagenomics (Fig. 2). To obtain CH_4_ oxidation rates at -20°C (1 × 10^-4^ nmol CH_4_ h^-1^ g^-1^), long-term Mackay Glacier microcosms from a previous study [2] were reanalysed after four years of continuous incubation (Fig. S8) and showed oxidation to below sub-atmospheric concentrations in three of the 12 microcosms. Mackay Glacier rates at -20°C were much lower than those measured at 4 and 10°C [2], though average cell-specific rates of CH_4_ oxidation at 4°C (1 × 10^-10^ ± 8 × 10^-11^) were similar to those for H_2_ (6 × 10^-10^ ± 3 × 10^-10^) and higher than CO (1 × 10^-11^ ± 2 × 10^-11^) across all nine soil samples reported here (Fig. 3BDF).

Temperature-rate relationships for each sample and region were modelled using General Temperature Dependence (GTD) theory (see Methods) to better understand the thermodynamics behind the wide thermal range of H_2_ and CO consumption. Rates were initially modelled with the Arrhenius equation, which describes a log-linear increase in reaction rate with temperature until enzyme denaturation or cell death, but here only fit the data from -20 to 42°C for each gas (Fig. S9, Table S6). GTD is based in Transition State Theory, which builds on Arrhenius to instead present a curved fit that is more applicable across the full range of the thermal sensitivity data [43]. Here, GTD predicted that soil samples from the colder, higher latitude sites in Dronning Maud Land had an average H_2_ oxidation *T*_opt_ of 13.3°C (9.3–17.9°C 95% CI), which is lower than soils from the milder regions of Robinson Ridge (*T*_opt_ 22.8°C; 19.1–27.4°C) and Bunger Hills (*T*_opt_ 18.4°C; 11.1–26.2°C) (Fig. 3, Table 1). GTD theory was also used to calculate the average ΔC for soil H_2_ oxidation across each region (Table 1). As demonstrated by other similar thermal sensitivity models based in transition state theory [83], this value can provide an indication of the overall flexibility of atmospheric trace gas oxidation potential in the ecosystem as the temperature changes [84]. The lower average ΔC value for Dronning Maud Land soils therefore suggests that they are better adapted to oxidising H_2_ at a wider range of temperatures, whereas those from Robinson Ridge and Bunger Hills seem better adapted to a warmer and narrower thermal range (Table 1). The thermodynamic flexibility of H_2_ oxidation and environmentally relevant *T*_opt_ support the foundational importance of this process in nutrient-poor Antarctic soils.

**Table 1.** Thermodynamic properties of H_2_ and CO oxidation in Antarctic soils, based on bulk soil oxidation rates. Values in square brackets indicate 95% confidence intervals based on 1000 bootstrap iterations of the model.

| Gas | Thermodynamic Parameter | All (*n*=9) | Bunger Hills (*n*=3) | Dronning Maud Land (*n*=3) | Robinson Ridge (*n*=3) |
| --- | --- | --- | --- | --- | --- |
| H<sub>2</sub> | *T*<sub>opt</sub> (°C) | 16.9 [13.8–20.7] | 18.4 [11.1–26.2] | 13.3 [9.3–17.9] | 22.8 [19.1–27.4] |
|  | Δ*C* (kJ mol<sup>-1</sup> K<sup>-1</sup>) | -3.40<br>[-4.61 to -2.66] | -2.74<br>[-5.29 to -1.72] | -3.77<br>[-6.07 to -2.54] | -2.94<br>[-3.64 to -2.01] |
|  | Δ*H* (kJ mol<sup>-1</sup>) | 986 [779–1330] | 799 [497–1530] | 1082<br>[725–1730] | 871 [609–1070] |
| CO | *T*<sub>opt</sub> (°C) | 25.3 [17.5–41.0] | 20.9 [14.9–39.2] | nd<sup>a</sup> | 21.7 [18.1–27.6] |
|  | Δ*C* (kJ mol<sup>-1</sup> K<sup>-1</sup>) | -3.10<br>[-4.67 to -1.56] | -5.21<br>[-10.40 to -1.26] | nd<sup>a</sup> | -4.20<br>[-5.16 to -2.62] |
|  | Δ*H* (kJ mol<sup>-1</sup>) | 926 [479–1360] | 1530<br>[381–3040] | nd<sup>a</sup> | 1240<br>[786–1510] |

CO oxidation is predicted to have higher temperature optima than for H_2_ (25.3°C compared to 16.9°C overall, Table 1, Fig. 3). Regionally, the Bunger Hills and Robinson Ridge soils peaked close together at 20.9°C and 21.7°C, respectively, (Table 1). Due to the far narrower temperature range over which CH_4_ consumption occurred, model fit to these data was considered unreliable and therefore not presented. For H_2_ and CO, however, model fits for bulk soil oxidation rates (iterated over 1000 bootstraps) were reasonable and had pseudo *R*^2^ values of 0.390 (H_2_) and 0.503 (CO), and Root Mean Squared Error (RMSE) values of 1.850 (H_2_) and 1.261 (CO). Model fits improved when considering the cell-specific oxidation rates (pseudo *R*^2^ values were 0.588 (H_2_) and 0.675 (CO); RMSE was 1.309 (H_2_) and 1.089 (CO)) overall supporting the generality of GTD, which had not previously been tested for any Antarctic system.

### Trace gas oxidation is spatially widespread and seasonally sustained in Antarctic ice-free regions

To further investigate the effect of temperature on trace gas oxidation and expand our predictions across the continent, generalised linear models (GLMs) with gamma error distributions were fitted to bulk H_2_, CO and CH_4_ oxidation rates (Fig. S10). Significant effects in the conditional component (i.e. using only non-zero rate values) indicate a temperature influence on oxidation rate, whereas significance in the zero-inflated model reflects a temperature effect on the binary presence or absence of activity. As expected, these models predicted that temperature significantly influences H_2_ and CO oxidation, with these processes peaking at 25°C and 37°C respectively. For both gases, the greatest changes were observed between ∼1–15°C and ∼31–46°C (Fig. S10A) and both linear and quadratic terms were significant in the conditional model, though effect sizes were small (Supplementary Text). Model prediction accuracy was better for CO than H_2_, with a much lower Mean Squared Error (1.79 × 10^-5^ compared to 2.07) and although temperature is significant, the effect size is small, likely a result of the small sample size. Nonetheless, a clear and strong effect of temperature was observed in the zero-inflation models for H_2_ and CO, i.e. on the presence/absence of activity (Supplementary Text). Temperature also had a significant effect on whether CH_4_ oxidation occurred but did not have a significant effect on rate (Fig. S10C). Given the absence of a rate effect for CH_4_, further analysis (continent-wide and future modelling) focused solely on H_2_ and CO.

The baseline relationship between temperature and H_2_ and CO oxidation rates was extrapolated to investigate putative rates across the entirety of ice-free Antarctica based on the average annual temperature per 1 km^2^ from 1981–2010 (Fig. 4AB). These values were then aggregated for each of the Antarctic Conservation Biogeographic Regions (ACBRs; Fig. 4CD). As expected, higher rates of H_2_ and CO oxidation are predicted in warmer, lower latitude regions, such as the South Orkney Islands and Antarctic Peninsula. Rates are predicted to be the lowest in the far colder mountainous or inland ACBRs (Fig. 4CD). Furthermore, oxidation rates in the current/near-future time period (2011-2040) exhibit a strong seasonality, with H_2_ and CO rates being 5.6-fold and 8.1-fold higher in the summer months compared to winter months, though importantly with significant rates still observed throughout winter (Fig. 5AB). It should be cautioned that these models solely consider temperature, yet other unaccounted factors (e.g. moisture, pH, micronutrients) are likely to also influence oxidation rates. Therefore, increasing the sample size, obtaining temperature-rate data from across all ACBRs, and considering additional parameters would further improve model accuracy, and is under consideration for future iterations. Regardless, the existing models strongly suggest that based on the thermal range of activities observed, trace gas oxidation can be sustained across ice-free Antarctica and throughout seasons.

**Figure 4.**
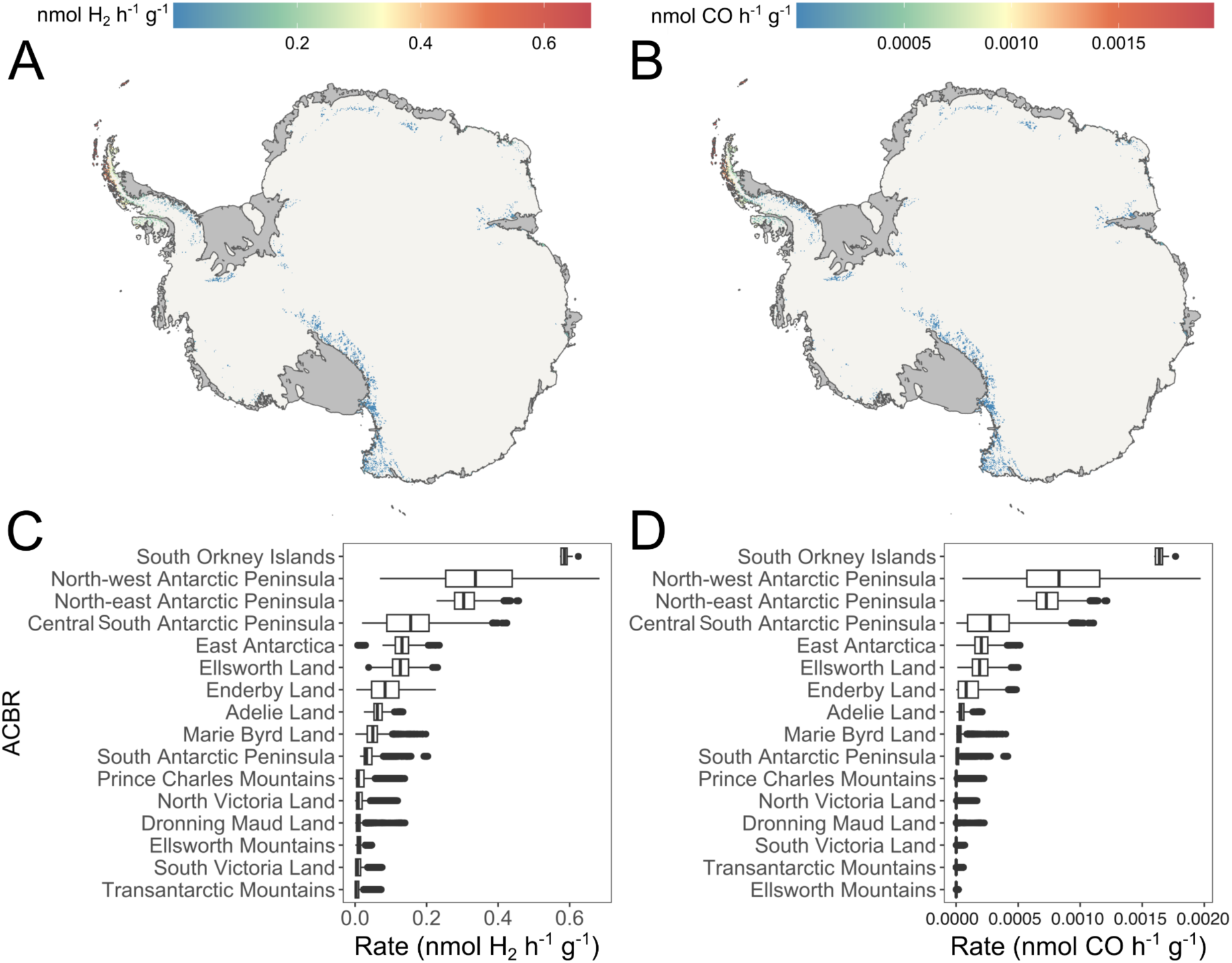
Predicted rates of H_2_ and CO oxidation across ice-free Antarctica. Map of Antarctica showing predicted baseline (1981–2010) oxidation rates for (**A**) H_2_ and (**B**) CO, averaged across an entire year. Bar charts show baseline annual average rates of (**C**) H_2_ and (**D**) CO oxidation across the ice-free regions broken down by Antarctic Conservation Bioregions (ACBRs), from highest to lowest rates. Note the different scales for H_2_ and CO oxidation in both the heatmaps and bar charts.

**Figure 5.**
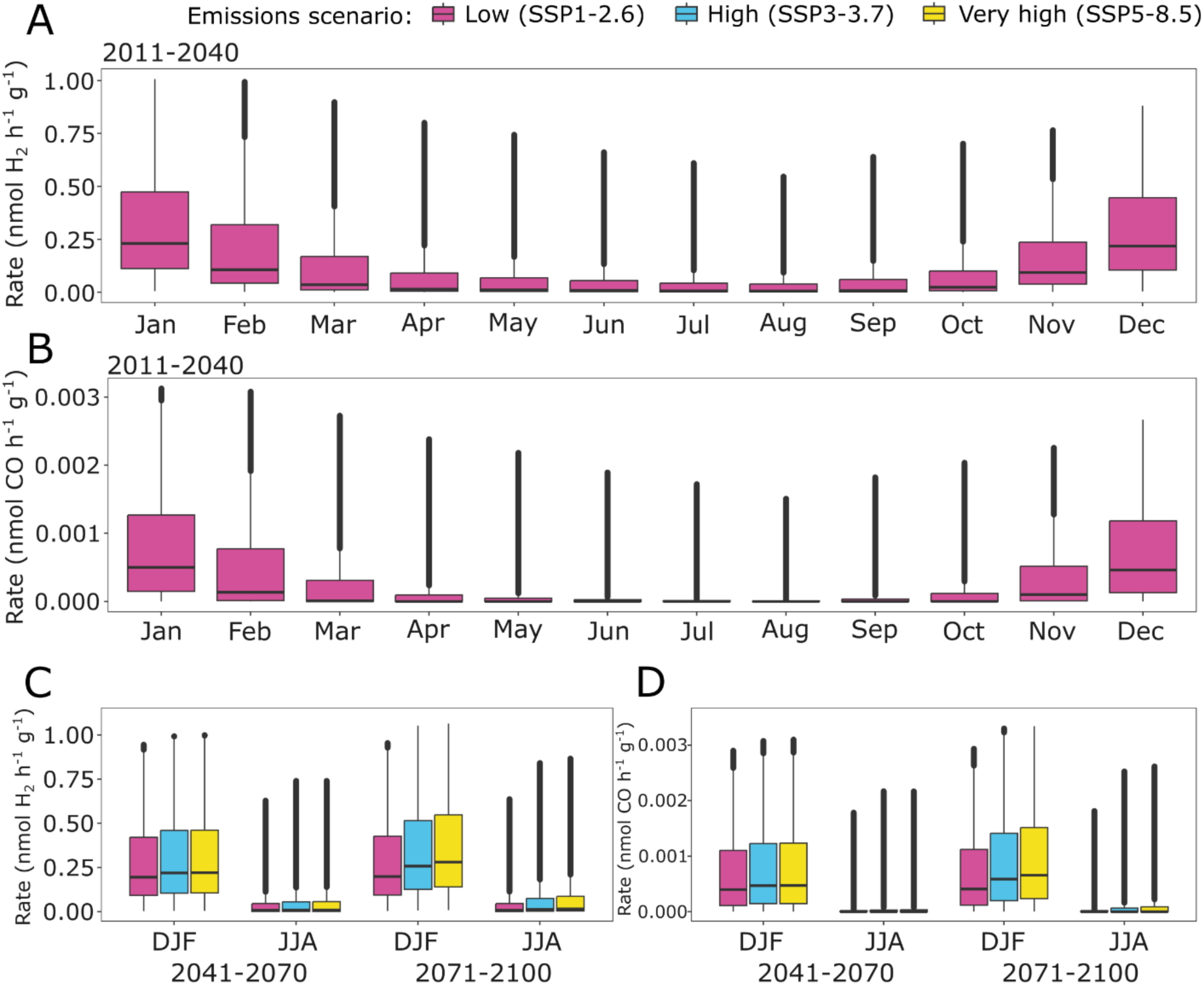
Variations in H_2_ and CO oxidation rates with changing seasons and emissions scenarios. Monthly (A) H_2_ and (B) CO oxidation rates over a 12-month period for the present and near future (2011–2040). Boxplots of (C) H_2_ and (D) CO oxidation rates in the austral summer (December, January, February) and winter (June, July, August) in the medium and far future, under three emission scenarios. Boxplots show median, interquartile range, minimum and maximum; all data points are included.

### Trace gas oxidation will respond to different warming scenarios

Earlier *ex situ* incubations suggested that trace gas oxidation rates will increase with the rising temperatures forecast due to anthropogenic climate warming. We used the oxidation-rate GLM to investigate this for H_2_ and CO oxidation rates across ice-free Antarctica under low (SSP1-2.6), high (SSP3-7.0), and very high (SSP5-8.5) emissions scenarios [85] in the mid-term (2041-2070) and long-term (2071–2100) future (Fig. 5). The strong seasonal variability in the current/near-future period (2011–2040) is expected to somewhat decline in future time periods, as winter (June, July, August) rates increase faster than summer (December, January, February) rates, especially under high and very high emissions. This follows a recent trend of warm extremes in winter being larger than those in summer [86]. By the end of the century, summer H_2_ oxidation rates may increase by 27% under high emissions or 35% under very high emissions (compared to SSP1-2.6 2011-2040; Fig. 5C). Similarly, summer CO oxidation rates may increase by 33% (high) to 44% (very high) (Fig. 5D). Differences between the SSPs widen over time: in the mid-term future (2041-2070), high or very high emissions could lead to a 9-10% increase from SSP1-2.6 summer rates. In the far future (2071-2100), this widens substantially to a 62-79% increase. Proportional increases in CO oxidation rates are similar, though may have less effect given the orders of magnitude lower rates overall (Table S7). In addition to increasing rates, ice-free regions of Antarctica are predicted to expand by up to 25% by 2100 [87] and as this happens the area available and hence volume of trace gas oxidation is likely to increase further.

## Discussion

Our findings demonstrate that trace gas oxidation is a widespread and temperature-resilient metabolic strategy in Antarctic soil ecosystems. Communities with similar composition, capabilities, and activities occupy soils from distant and contrasting regions: the sharp mountains and nunataks of Dronning Maud Land, and coastal oases in Bunger Hills, Robinson Ridge, and Schirmacher Oasis. In each case, the dominant community members are aerobic bacteria (especially Actinomycetota) that use atmospheric trace gases to meet energy and carbon needs amid organic carbon starvation. These communities are remarkably compositionally and functionally similar to those reported from Victoria Land, the Transantarctic Mountains, the Antarctic Peninsula, and the Windmill Islands [1, 2, 88], as well as some endolithic communities [89], and trace gas oxidation is likely to be the dominant process sustaining them in all cases. Altogether, *in situ* observations, *ex situ* incubations, and rate-temperature modelling suggest that atmospheric H_2_ oxidation is particularly widespread, with this gas likely to be consumed at considerable rates at subzero temperatures during austral winters and also at elevated temperatures far beyond Antarctic norms. Consistently, both genomic and biogeochemical analyses suggest that Antarctic microbes generally function far below their temperature optima. This finding aligns with previous studies demonstrating that many microbes found at low temperatures tend to have optimal growth temperatures substantially higher than their natural environment [90], including a number of Antarctic isolates (e.g. [36, 91]). This thermal breadth, underpinned by the widespread genetic capacity for aerotrophy across multiple bacterial phyla, highlights Antarctic microbes as both resilient and resourceful in extreme environments.

These results extend understanding of both the upper and lower thermal limits for metabolic activity in Antarctica. We observed an upper functional (or survival) limit of 75°C for H_2_ oxidation, which far exceeds the temperatures that Antarctic microbes are ever likely to naturally experience, except in an isolated geothermally active region in Victoria Land [92]. Although some thermotolerant microbes may spread from this area, dispersal within and into the continent is considered highly restricted, due to the vast distances between ice-free refugia [93]. Conversely, the subzero temperature range investigated here demonstrates that Antarctic trace gas oxidisers can produce energy throughout the winter months, given that atmospheric gas diffusion can still occur even amid increased snow cover [94]. H_2_ oxidation still occurred rapidly at -20°C, within the proposed lower limit for cellular metabolism (-10 to -26°C) where cells are thought to undergo vitrification [95], though in line with several Arctic studies reporting activities even beyond this range [96–99]. Further work should determine the lowest limits of trace gas oxidation activity and explore which other cellular activities occur at such temperatures. The 95°C range of H_2_ oxidation reported in this work builds upon enzyme activity and soil respiration measurements from 4-70°C in a tropical soil [100] and chemoautotrophy observed in Yellowstone Lake thermal vent waters from 15-80°C [101]. We expect that the key taxa mediating trace gas oxidation in the Antarctic soils reported here vary between temperatures, similar to what has been shown in Alaskan permafrost soils [96]. The widespread presence of Group 1 [NiFe]-hydrogenases across 11 phyla (Fig. 2A, Table S4) provides high functional redundancy for this trait and likely contributes to its thermal resilience.

Rate-temperature response curves also demonstrated that diverse Antarctic soils conform to a previously asserted universal general model for temperature dependence [43]. To date, this “universality” had not been tested for any ecosystem functions across the Antarctic continent, which is often defined by its unique biodiversity and features. Nonetheless, despite a wide thermal breadth for H_2_ and CO oxidation rates, the thermodynamic parameter -ΔC values were within the most commonly found range for biological processes. Likewise, optimum temperatures were typical of or close to those for most systems (∼25°C) [43]. Thus, despite trace gas oxidation being a process quite different from others examined at the molecular level for generality, the general theory for temperature dependence holds up well for it, providing additional support for this idea. More broadly, the thermal sensitivity modelling for trace gas oxidation within Antarctic soils enables a more comprehensive understanding of terrestrial ecosystem functioning across the continent and supports projections of soil ecosystem functions under current and future climates.

Although H_2_ and CO oxidation are thermally robust, our results suggest their mediators still respond to environmental conditions. Temperature clearly constrains the rates of their oxidation and warming could relax these constraints, expanding the realised niche of trace gas oxidisers and potentially enhancing their ecological role. Our modelling predicts increases in activity with rising temperatures (up to ∼37°C), especially in colder inland regions and during winter months, indicating that climate change may amplify the energy flux through these pathways. This complements previous observations of increased respiration in Antarctic Peninsula soils under experimental warming at temperatures up to 50°C [39], and reports of increased methane oxidation in experimentally warmed Arctic permafrost [102–104]. In contrast, CH_4_ oxidation in Antarctic soils occurs less consistently and within narrower thermal ranges, with the few mediators of this process much more sensitive to short- and long-term variations in environmental conditions. Although rates of CH_4_ oxidation are likely to increase in the future, it seems unlikely that this would occur at a scale to mitigate other contributions to climate forcing globally.

Nonetheless, the future trajectory of Antarctica’s aerotrophic communities is uncertain. As the continent warms and conditions become increasingly favourable for photosynthetic microbes and pioneer plants, trace gas oxidisers may face intensified competition for space, nutrients, and possibly even redox niches. Whether aerotrophs will persist as foundational primary producers, be displaced, or shift to new ecological roles remains a critical open question. Our study provides strong evidence for the current dominance and future resilience of trace gas oxidation, but our understanding remains incomplete. Temperature is only one of many environmental variables shaping microbial community composition and activity. Climate change is also causing increased precipitation over certain Antarctic regions, with positive or negative effects on photosynthetic vegetation dependent on location [105–107], and growing plant populations will increase nutrient inputs to the soil [108]. Expanding studies to include broader geographic regions, soil physicochemical parameters, and seasonal sampling will be essential to fully unravel the factors that determine microbial distributions and interactions in Antarctic soils. These insights will be vital for predicting how life in Earth’s coldest deserts may respond to rapid environmental change, and to what extent the processes and mediators of primary production change.

## End Matter

### Author Contributions

CG, SIH, TFH, PML, SKB, MAM, and SLC conceived and designed the work. SIH, DAC, RL, WPAL, TT and AB carried out field work. TFH, SIH, RL, TJ, and WPAL performed the experiments. DAC and SIH performed modelling. SIH, TFH, DAC, FR and CG analysed the data. SIH, TFH, CG, and DAC wrote the manuscript. All authors edited and approved the final manuscript.

## Acknowledgements

The Australian Antarctic Division is thanked for field support through AAS Project 4628, for collection of the Robinson Ridge and Bunger Hills samples. We are grateful to White Desert for field and logistical support on the Dronning Maud Land expeditions. Thank you to Ms Laura M. Phillips for assistance collecting soils from the Bunger Hills and to Ms Noémie L.M. Sheppard for her assistance with R coding. This study used the MASSIVE M3 supercomputing infrastructure. This work contributes to delivering the Australian Antarctic Science Decadal Strategy.

## Data Availability

Metagenomic reads are available on the NCBI Sequence Read Archive under PRJNA1272669 and MAGs are available on NCBI under BioProject PRJNA1272656 and PRJNA1272657. Code for the general temperature dependence theory fit is available at: https://github.com/micro-ry/Antarctic-soils-thermal-sensitivity. Code for the continent-wide and future oxidation rate models is available at https://github.com/DavidAClarke/antarctic-trace-gas-modelling.

## References

1. Ji M, Greening C, Vanwonterghem I, et al. Atmospheric trace gases support primary production in Antarctic desert surface soil. Nature 2017;552:400–403. 10.1038/nature25014

2. Ortiz M, Leung PM, Shelley G, et al. Multiple energy sources and metabolic strategies sustain microbial diversity in Antarctic desert soils. Proc Natl Acad Sci 2021;118:e2025322118. 10.1073/pnas.2025322118

3. Bockheim JG. Properties and classification of cold desert soils from Antarctica. Soil Sci Soc Am J 1997;61:224–231. 10.2136/sssaj1997.03615995006100010031x

4. Beyer L, Bölter M (eds). Geoecology of Antarctic Ice-Free Coastal Landscapes, 1st edn. Springer Berlin, Heidelberg, 2002.

5. Pointing SB, Chan Y, Lacap DC, et al. Highly specialized microbial diversity in hyper-arid polar desert. Proc Natl Acad Sci 2009;106:19964–19969. 10.1073/pnas.0908274106

6. Lee CK, Barbier BA, Bottos EM, et al. The Inter-Valley Soil Comparative Survey: the ecology of Dry Valley edaphic microbial communities. ISME J 2012;6:1046–1057. 10.1038/ismej.2011.170

7. Lebre PH, Bosch J, Coclet C, et al. Expanding Antarctic biogeography: microbial ecology of Antarctic island soils. Ecography 2023;e06568. 10.1111/ecog.06568

8. Harris JM, Tibbles BJ. Factors affecting bacterial productivity in soils on isolated inland nunataks in continental Antarctica. Microb Ecol 1997;33:106–123. 10.1007/s002489900013

9. Cary SC, McDonald IR, Barrett JE, et al. On the rocks: the microbiology of Antarctic Dry Valley soils. Nat Rev Microbiol 2010;8:129–138. 10.1038/nrmicro2281

10. Dragone NB, Diaz MA, Hogg ID, et al. Exploring the boundaries of microbial habitability in soil. J Geophys Res Biogeosciences 2021;126:e2020JG006052. 10.1029/2020JG006052

11. Wood C, Bruinink A, Trembath-Reichert E, et al. Active microbiota persist in dry permafrost and active layer from Elephant Head, Antarctica. ISME Commun 2024;4. 10.1093/ismeco/ycad002

12. Ray AE, Zhang E, Terauds A, et al. Soil microbiomes with the genetic capacity for atmospheric chemosynthesis are widespread across the poles and are associated with moisture, carbon, and nitrogen limitation. Front Microbiol 2020;11:1936. 10.3389/fmicb.2020.01936

13. Ray AE, Zaugg J, Benaud N, et al. Atmospheric chemosynthesis is phylogenetically and geographically widespread and contributes significantly to carbon fixation throughout cold deserts. ISME J 2022;16:2547–2560. 10.1038/s41396-022-01298-5

14. Bay S, Ni G, Lappan R, et al. Microbial aerotrophy enables continuous primary production in diverse cave ecosystems. Nat Commun 2025;**Accepted in principle**.

15. Bay SK, Dong X, Bradley JA, et al. Trace gas oxidizers are widespread and active members of soil microbial communities. Nat Microbiol 2021;6:246–256. 10.1038/s41564-020-00811-w

16. Jordaan K, Lappan R, Dong X, et al. Hydrogen-oxidizing bacteria are abundant in desert soils and strongly stimulated by hydration. mSystems 2020;5:e01131–20. 10.1128/mSystems.01131-20

17. Liu L, Chen Y, Shen J, et al. Metabolic versatility of soil microbial communities below the rocks of the hyperarid Dalangtan Playa. Appl Environ Microbiol 2023;89:e01072–23. 10.1128/aem.01072-23

18. Lappan R, Shelley G, Islam ZF, et al. Molecular hydrogen in seawater supports growth of diverse marine bacteria. Nat Microbiol 2023;8:581–595. 10.1038/s41564-023-01322-0

19. Kotzé C, Meiklejohn I. Temporal variability of ground thermal regimes on the northern buttress of the Vesleskarvet nunatak, western Dronning Maud Land, Antarctica. Antarct Sci 2017;29:73–81. 10.1017/S095410201600047X

20. 20. Environmental Management Unit. Davis Aerodrome Project (DAP) – Subsurface Soil Temperature Monitoring Data. 2024. Australian Antarctic Data Centre, 2024.

21. Convey P, Coulson SJ, Worland MR, et al. The importance of understanding annual and shorter-term temperature patterns and variation in the surface levels of polar soils for terrestrial biota. Polar Biol 2018;41:1587–1605. 10.1007/s00300-018-2299-0

22. Thompson DC, Bromley AM, Craig RMF. Ground temperatures in an Antarctic dry valley. N Z J Geol Geophys 1971;14:477–483. 10.1080/00288306.1971.10421941

23. Prabhu Matondkar SG. Third Indian Expedition to Antarctica, Scientific Report, 1986. Department of Ocean Development, 1986.

24. Sedov S, Zazovskaya E, Fedorov-Davydov D, et al. Soils of East Antarctic oasis: interplay of organisms and mineral components at microscale. Bol Soc Geológica Mex 2019;71:43–63. 10.18268/BSGM2019v71n1a4

25. Chown SL, Leihy RI, Naish TR, et al. Antarctic Climate Change and the Environment: A Decadal Synopsis and Recommendations for Action. Cambridge, United Kingdom: Scientific Committee on Antarctic Research, 2022.

26. González-Herrero S, Barriopedro D, Trigo RM, et al. Climate warming amplified the 2020 record-breaking heatwave in the Antarctic Peninsula. Commun Earth Environ 2022;3:122. 10.1038/s43247-022-00450-5

27. Robinson SA, Klekociuk AR, King DH, et al. The 2019/2020 summer of Antarctic heatwaves. Glob Change Biol 2020;26:3178–3180. 10.1111/gcb.15083

28. Wille JD, Alexander SP, Amory C, et al. The extraordinary March 2022 East Antarctica “heat” wave. Part I: Observations and meteorological drivers. J Clim 2024;37:757–778. 10.1175/JCLI-D-23-0175.1

29. Onofri S, Fenice M, Cicalini AR, et al. Ecology and biology of microfungi from Antarctic rocks and soils. Ital J Zool 2000;67:163–167. 10.1080/11250000009356372

30. Kim D, Park HJ, Kim JH, et al. Passive warming effect on soil microbial community and humic substance degradation in maritime Antarctic region. J Basic Microbiol 2018;58:513–522. 10.1002/jobm.201700470

31. Newsham KK, Tripathi BM, Dong K, et al. Bacterial community composition and diversity respond to nutrient amendment but not warming in a maritime Antarctic soil. Microb Ecol 2019;78:974–984. 10.1007/s00248-019-01373-z

32. Newsham KK, Misiak M, Goodall-Copestake WP, et al. Experimental warming increases fungal alpha diversity in an oligotrophic maritime Antarctic soil. Front Microbiol 2022;13:1050372. 10.3389/fmicb.2022.1050372

33. Wang C, Morrissey EM, Mau RL, et al. The temperature sensitivity of soil: microbial biodiversity, growth, and carbon mineralization. ISME J 2021;15:2738–2747. 10.1038/s41396-021-00959-1

34. Sáez-Sandino T, García-Palacios P, Maestre FT, et al. The soil microbiome governs the response of microbial respiration to warming across the globe. Nat Clim Change 2023;13:1382–1387. 10.1038/s41558-023-01868-1

35. Cruz-Paredes C, Tájmel D, Rousk J. Variation in temperature dependences across Europe reveals the climate sensitivity of soil microbial decomposers. Appl Environ Microbiol 2023;89:e02090–22. 10.1128/aem.02090-22

36. Valenzuela-Ibaceta F, Carrasco V, Lagos-Moraga S, et al. *Arthrobacter vasquezii* sp. nov., isolated from a soil sample from Union Glacier, Antarctica. Int J Syst Evol Microbiol 2023;73. 10.1099/ijsem.0.006095

37. Vodickova P, Suman J, Benesova E, et al. *Arthrobacter polaris* sp. nov., a new cold-adapted member of the family Micrococcaceae isolated from Antarctic fellfield soil. Int J Syst Evol Microbiol 2022;72. 10.1099/ijsem.0.005541

38. Ferguson SH, Powell SM, Snape I, et al. Effect of temperature on the microbial ecology of a hydrocarbon-contaminated Antarctic soil: Implications for high temperature remediation. Cold Reg Sci Technol 2008;53:115–129. 10.1016/j.coldregions.2007.04.006

39. Laudicina VA, Benhua S, Dennis PG, et al. Responses to increases in temperature of heterotrophic micro-organisms in soils from the maritime Antarctic. Polar Biol 2015;38:1153– 1160. 10.1007/s00300-015-1673-4

40. De Souza Carvalho JV, De Sá Mendonça E, Barbosa RT, et al. Impact of expected global warming on C mineralization in maritime Antarctic soils: Results of laboratory experiments. Antarct Sci 2010;22:485–493. 10.1017/S0954102010000258

41. Bokhorst S, Huiskes A, Convey P, et al. Climate change effects on organic matter decomposition rates in ecosystems from the Maritime Antarctic and Falkland Islands. Glob Change Biol 2007;13:2642–2653. 10.1111/j.1365-2486.2007.01468.x

42. Yergeau E, Kowalchuk GA. Responses of Antarctic soil microbial communities and associated functions to temperature and freeze-thaw cycle frequency. Environ Microbiol 2008;10:2223–2235. 10.1111/j.1462-2920.2008.01644.x

43. Arroyo JI, Díez B, Kempes CP, et al. A general theory for temperature dependence in biology. Proc Natl Acad Sci 2022;119:e2119872119. 10.1073/pnas.2119872119

44. Ohtani S, Kanda H. Dronning Maud Land and Its Environments. In: Beyer L, Bölter M (eds), Geoecology of Antarctic Ice-Free Coastal Landscapes. Berlin, Heidelberg: Springer Berlin Heidelberg, 2002, 51–68.

45. Alberts F, Blodgett G. The Bunger Hills area of Antarctica. Prof Geogr 1956;8:13–15. 10.1111/j.0033-0124.1956.83_13.x

46. Salazar VW, Shaban B, Quiroga MDM, et al. Metaphor—A workflow for streamlined assembly and binning of metagenomes. GigaScience 2022;12:giad055. 10.1093/gigascience/giad055

47. Chen S. Ultrafast one-pass FASTQ data preprocessing, quality control, and deduplication using fastp. iMeta 2023;2:e107. 10.1002/imt2.107

48. Li D, Liu C-M, Luo R, et al. MEGAHIT: an ultra-fast single-node solution for large and complex metagenomics assembly via succinct *de Bruijn* graph. Bioinformatics 2015;31:1674–1676. 10.1093/bioinformatics/btv033

49. Gruber-Vodicka HR, Seah BKB, Pruesse E. phyloFlash: Rapid small-subunit rRNA profiling and targeted assembly from metagenomes. mSystems 2020;5:e00920–20. 10.1128/mSystems.00920-20

50. Nissen JN, Johansen J, Allesøe RL, et al. Improved metagenome binning and assembly using deep variational autoencoders. Nat Biotechnol 2021;39:555–560. 10.1038/s41587-020-00777-4

51. Kang DD, Li F, Kirton E, et al. MetaBAT 2: an adaptive binning algorithm for robust and efficient genome reconstruction from metagenome assemblies. PeerJ 2019;7:e7359. 10.7717/peerj.7359

52. Pan S, Zhao X-M, Coelho LP. SemiBin2: self-supervised contrastive learning leads to better MAGs for short- and long-read sequencing. Bioinformatics 2023;39:i21–i29. 10.1093/bioinformatics/btad209

53. Alneberg J, Bjarnason BS, De Bruijn I, et al. Binning metagenomic contigs by coverage and composition. Nat Methods 2014;11:1144–1146. 10.1038/nmeth.3103

54. Uritskiy GV, DiRuggiero J, Taylor J. MetaWRAP—a flexible pipeline for genome-resolved metagenomic data analysis. Microbiome 2018;6:158. 10.1186/s40168-018-0541-1

55. Olm MR, Brown CT, Brooks B, et al. dRep: a tool for fast and accurate genomic comparisons that enables improved genome recovery from metagenomes through de-replication. ISME J 2017;11:2864–2868. 10.1038/ismej.2017.126

56. Chklovski A, Parks DH, Woodcroft BJ, et al. CheckM2: a rapid, scalable and accurate tool for assessing microbial genome quality using machine learning. Nat Methods 2023;20:1203– 1212. 10.1038/s41592-023-01940-w

57. The Genome Standards Consortium, Bowers RM, Kyrpides NC, et al. Minimum information about a single amplified genome (MISAG) and a metagenome-assembled genome (MIMAG) of bacteria and archaea. Nat Biotechnol 2017;35:725–731. 10.1038/nbt.3893

58. Chaumeil P-A, Mussig AJ, Hugenholtz P, et al. GTDB-Tk v2: memory friendly classification with the genome taxonomy database. Bioinformatics 2022;38:5315–5316. 10.1093/bioinformatics/btac672

59. Parks DH, Chuvochina M, Rinke C, et al. GTDB: an ongoing census of bacterial and archaeal diversity through a phylogenetically consistent, rank normalized and complete genome-based taxonomy. Nucleic Acids Res 2022;50:D785–D794. 10.1093/nar/gkab776

60. Aroney STN, Newell RJP, Nissen JN, et al. CoverM: read alignment statistics for metagenomics. Bioinformatics 2025;41:btaf147. 10.1093/bioinformatics/btaf147

61. Kalyaanamoorthy S, Minh BQ, Wong TKF, et al. ModelFinder: fast model selection for accurate phylogenetic estimates. Nat Methods 2017;14:587–589. 10.1038/nmeth.4285

62. Minh BQ, Schmidt HA, Chernomor O, et al. IQ-TREE 2: New Models and Efficient Methods for Phylogenetic Inference in the Genomic Era. Mol Biol Evol 2020;37:1530–1534. 10.1093/molbev/msaa015

63. Hoang DT, Chernomor O, Von Haeseler A, et al. UFBoot2: Improving the Ultrafast Bootstrap Approximation. Mol Biol Evol 2018;35:518–522. 10.1093/molbev/msx281

64. Letunic I, Bork P. Interactive Tree Of Life (iTOL): an online tool for phylogenetic tree display and annotation. Bioinformatics 2007;23:127–128. 10.1093/bioinformatics/btl529

65. Hyatt D, Chen G-L, LoCascio PF, et al. Prodigal: prokaryotic gene recognition and translation initiation site identification. BMC Bioinformatics 2010;11:119. 10.1186/1471-2105-11-119

66. Altschul S. Gapped BLAST and PSI-BLAST: a new generation of protein database search programs. Nucleic Acids Res 1997;25:3389–3402. 10.1093/nar/25.17.3389

67. Leung PM, Greening C. Greening lab metabolic marker gene databases. 2020. Monash University Collection, 2020.

68. Li G, Rabe KS, Nielsen J, et al. Machine learning applied to predicting microorganism growth temperatures and enzyme catalytic optima. ACS Synth Biol 2019;8:1411–1420. 10.1021/acssynbio.9b00099

69. Buchfink B, Xie C, Huson DH. Fast and sensitive protein alignment using DIAMOND. Nat Methods 2015;12:59–60. 10.1038/nmeth.3176

70. Nauer PA, Chiri E, Jirapanjawat T, et al. Technical note: Inexpensive modification of exetainers for the reliable storage of trace-level hydrogen and carbon monoxide gas samples. Biogeosciences 2021;18:729–737. 10.5194/bg-18-729-2021

71. Islam ZF, Cordero PRF, Feng J, et al. Two Chloroflexi classes independently evolved the ability to persist on atmospheric hydrogen and carbon monoxide. ISME J 2019;13:1801– 1813. 10.1038/s41396-019-0393-0

72. Brooks M E, Kristensen K, Benthem K J,van, et al. glmmTMB balances speed and flexibility among packages for zero-inflated generalized linear mixed modeling. R J 2017;9:378. 10.32614/RJ-2017-066

73. Hartig F. DHARMa: Residual Diagnostics for Hierarchical (Multi-Level / Mixed) Regression Models. 2024. 2024.

74. Fox J, Monette G. cv: Cross-Validating Regression Models. 2025. 2025.

75. Karger DN, Conrad O, Böhner J, et al. Climatologies at high resolution for the earth’s land surface areas. Sci Data 2017;4:170122. 10.1038/sdata.2017.122

76. Karger DN, Conrad O, Böhner J, et al. Climatologies at high resolution for the earth’s land surface areas. 2021. EnviDat, 2021.

77. Terauds A, Chown SL, Morgan F, et al. Conservation biogeography of the Antarctic. Divers Distrib 2012;18:726–741. 10.1111/j.1472-4642.2012.00925.x

78. Terauds A, Lee JR. Antarctic biogeography revisited: updating the Antarctic Conservation Biogeographic Regions. Divers Distrib 2016;22:836–840. 10.1111/ddi.12453

79. Miyazaki M, Sakai S, Yamanaka Y, et al. *Spirochaeta psychrophila* sp. nov., a psychrophilic spirochaete isolated from subseafloor sediment, and emended description of the genus Spirochaeta. Int J Syst Evol Microbiol 2014;64:2798–2804. 10.1099/ijs.0.062463-0

80. Tahon G, Tytgat B, Lebbe L, et al. *Abditibacterium utsteinense* sp. nov., the first cultivated member of candidate phylum FBP, isolated from ice-free Antarctic soil samples. Syst Appl Microbiol 2018;41:279–290. 10.1016/j.syapm.2018.01.009

81. Labeda DP. *Crossiella* gen. nov., a new genus related to *Streptoalloteichus*. Int J Syst Evol Microbiol 2001;51:1575–1579. 10.1099/00207713-51-4-1575

82. Leung PM. Energetic Basis of Microbial Growth and Persistence in Terrestrial Oligotrophic Ecosystems. 2023. Doctorate, Monash University.

83. Hobbs JK, Jiao W, Easter AD, et al. Change in heat capacity for enzyme catalysis determines temperature dependence of enzyme catalyzed rates. ACS Chem Biol 2013;8:2388–2393. 10.1021/cb4005029

84. Schipper LA, Hobbs JK, Rutledge S, et al. Thermodynamic theory explains the temperature optima of soil microbial processes and high *Q*_10_ values at low temperatures. Glob Change Biol 2014;20:3578–3586. 10.1111/gcb.12596

85. IPCC. Summary for Policymakers. In: Masson-Delmotte V, Zhai P, Pirani A, et al. (eds), Climate Change 2021 – The Physical Science Basis. Contribution of Working Group I to the Sixth Assessment Report of the Intergovernmental Panel on Climate Change, 1st edn. Cambridge University Press, 2023, 3–32.

86. Xu M, Pithan F, Yang Q. Antarctic warm extremes across seasons and their response to advection. J Geophys Res Atmospheres 2024;129:e2024JD040884. 10.1029/2024JD040884

87. Lee JR, Raymond B, Bracegirdle TJ, et al. Climate change drives expansion of Antarctic ice-free habitat. Nature 2017;547:49–54. 10.1038/nature22996

88. Dragone NB, Childress MK, Vanderburgh C, et al. A comprehensive survey of soil microbial diversity across the Antarctic continent. Polar Biol 2025;48:50. 10.1007/s00300-025-03372-y

89. Coleine C, Albanese D, Ray AE, et al. Metagenomics untangles potential adaptations of Antarctic endolithic bacteria at the fringe of habitability. Sci Total Environ 2024;917:170290. 10.1016/j.scitotenv.2024.170290

90. Knoll AH, Bauld J. The evolution of ecological tolerance in prokaryotes. Earth Environ Sci Trans R Soc Edinb 1989;80:209–223. 10.1017/S0263593300028650

91. Muñoz-Villagrán C, Grossolli-Gálvez J, Acevedo-Arbunic J, et al. Characterization and genomic analysis of two novel psychrotolerant *Acidithiobacillus ferrooxidans* strains from polar and subpolar environments. Front Microbiol 2022;13. 10.3389/fmicb.2022.960324

92. Bottos EM, Scarrow JW, Archer SDJ, et al. Bacterial Community Structures of Antarctic Soils. In: Cowan DA (ed.), Antarctic Terrestrial Microbiology. Berlin, Heidelberg: Springer, 2014.

93. McGeoch MA, Lee JR, Affleck S, et al. Ecological processes shaping Antarctic terrestrial biodiversity change. Nat Rev Biodivers ;Accepted.

94. Sommerfeld RA, Mosier AR, Musselman RC. CO_2_, CH_4_ and N_2_O flux through a Wyoming snowpack and implications for global budgets. Nature 1993;361. 10.1038/361140a0

95. Clarke A, Morris GJ, Fonseca F, et al. A low temperature limit for life on Earth. PLoS ONE 2013;8:e66207. 10.1371/journal.pone.0066207

96. Tuorto SJ, Darias P, McGuinness LR, et al. Bacterial genome replication at subzero temperatures in permafrost. ISME J 2014;8:139–149. 10.1038/ismej.2013.140

97. Rivkina EM, Friedmann EI, McKay CP, et al. Metabolic activity of permafrost bacteria below the freezing point. Appl Environ Microbiol 2000;66:3230–3233. 10.1128/AEM.66.8.3230-3233.2000

98. Elberling B, Brandt KK. Uncoupling of microbial CO_2_ production and release in frozen soil and its implications for field studies of arctic C cycling. Soil Biol Biochem 2003;35:263–272. 10.1016/S0038-0717(02)00258-4

99. Panikov NS, Flanagan PW, Oechel WC, et al. Microbial activity in soils frozen to below - 39°C. Soil Biol Biochem 2006;38:785–794. 10.1016/j.soilbio.2005.07.004

100. Paz-Ferreiro J, Fu S, Méndez A, et al. Biochar modifies the thermodynamic parameters of soil enzyme activity in a tropical soil. J Soils Sediments 2015;15:578–583. 10.1007/s11368-014-1029-7

101. Yang T, Lyons S, Aguilar C, et al. Microbial communities and chemosynthesis in Yellowstone Lake sublacustrine hydrothermal vent waters. Front Microbiol 2011;2. 10.3389/fmicb.2011.00130

102. Knoblauch C, Beer C, Liebner S, et al. Methane production as key to the greenhouse gas budget of thawing permafrost. Nat Clim Change 2018;8:309–312. 10.1038/s41558-018-0095-z

103. Zheng J, RoyChowdhury T, Yang Z, et al. Impacts of temperature and soil characteristics on methane production and oxidation in Arctic tundra. Biogeosciences 2018;15:6621–6635. 10.5194/bg-15-6621-2018

104. Oh Y, Zhuang Q, Liu L, et al. Reduced net methane emissions due to microbial methane oxidation in a warmer Arctic. Nat Clim Change 2020;10:317–321. 10.1038/s41558-020-0734-z

105. Robinson SA, King DH, Bramley-Alves J, et al. Rapid change in East Antarctic terrestrial vegetation in response to regional drying. Nat Clim Change 2018;8:879–884. 10.1038/s41558-018-0280-0

106. Roland TP, Bartlett OT, Charman DJ, et al. Sustained greening of the Antarctic Peninsula observed from satellites. Nat Geosci 2024;17:1121–1126. 10.1038/s41561-024-01564-5

107. Cannone N, Guglielmin M, Malfasi F, et al. Rapid soil and vegetation changes at regional scale in continental Antarctica. Geoderma 2021;394:115017. 10.1016/j.geoderma.2021.115017

108. Hill PW, Farrar J, Roberts P, et al. Vascular plant success in a warming Antarctic may be due to efficient nitrogen acquisition. Nat Clim Change 2011;1:50–53. 10.1038/nclimate1060

