## Supplementary Information and Figures for "Resilient Antarctic soil bacteria consume trace gases across wide temperature ranges"

#### Supplementary Text

**Generalised linear models to predict trace gas oxidation rates from temperature data**

To investigate the effect of temperature on rates of trace gas oxidation, zero-inflated generalised linear models (GLM) with gamma error distributions were fitted to bulk $H_2$, CO and $CH_4$ oxidation rates (Fig. S5). There was a significant effect of temperature on rates of $H_2$ oxidation, in both the conditional and zero-inflated components of the model. Hydrogen oxidation rates ranged from 0.0001 to 8.65 nmol $H_2$ $h^{-1}$ $g^{-1}$ between -20 to 75 ˚C; with the predicted peak rate occurring at 25°C (Fig. S10A). The greatest changes in the rate of hydrogen oxidation occurred over the temperature ranges of 1 - 13 °C and 35 - 46 °C. For the conditional model, both the linear and quadratic terms were significant, though effects were small with temperature estimates of $e^{-11.0894}$ = 1.53 x $10^{-5}$ [95% CI 2.32 x $10^{-7}$, 1.00 x $10^{-3}$] and $e^{-19.3284}$ = 4.03 x $10^{-9}$ [95% CI 9.00 x $10^{-11}$, 1.80 x $10^{-7}$] for the first and second-order term, respectively. Effects were stronger in the zero-inflation model, with temperature exerting a significant effect on the presence of hydrogen oxidation, with estimates of $e^{25.6874}$ = 1.43 x $10^{11}$ [95% CI 9.73 x $10^{6}$, 2.10 x $10^{15}$] and $e^{20.7044}$ = 9.81 x $10^{8}$ [95% CI 2.97 x $10^{4}$, 3.25 x $10^{13}$] for the linear and quadratic terms, respectively.

Rates of CO oxidation were also significantly affected by temperature. However, the linear term of the conditional model was not significant. Model prediction accuracy had a much lower mean squared error (MSE) of 1.79 x $10^{-5}$ compared to 2.07 for $H_2$. Oxidation rates were also lower than those for $H_2$, ranging from 2 x $10^{-5}$ to 0.028 nmol CO $hr^{-1}$ $g^{-1}$ (Fig. S10B). The peak rate of 0.028 nmol CO $h^{-1}$ $g^{-1}$ occurred at a greater temperature (37°C) than $H_2$. The greatest changes in the rate of carbon monoxide oxidation occurred over the temperature ranges of 4-15°C and 31-43°C. Effects of temperature on rates of CO oxidation were again small, with temperature estimates of $e^{-8.7058}$ = 1.66 x $10^{-4}$ [95% CI 1.68 x $10^{-10}$, 1.63 x $10^{2}$] and $e^{-20.6470}$ = 1.08 x $10^{-9}$ [95% CI 3.59 x $10^{-14}$, 3.25 x $10^{-5}$] for the first and second-order term, respectively. There is a very clear and strong effect of temperature in the zero-inflation model, with estimates of $e^{85.2661}$ = 1.07 x $10^{37}$ [95% CI 2.05 x $10^{21}$, 5.61 x $10^{52}$] and $e^{60.4947}$ = 1.87 x $10^{26}$ [95% CI 4.56 x $10^{15}$, 7.68 x $10^{36}$] for the linear and quadratic terms, respectively.

Unlike rates of hydrogen and carbon monoxide, there was no significant effect of temperature on rates of methane oxidation for neither term of the conditional model. Rates of methane oxidation range from 1.8 x $10^{-4}$ to 0.06 nmol $CH_4$ $h^{-1}$ $g^{-1}$, however, there is no clear relationship with temperature (Fig. S10C), other than the significant linear and quadratic terms of the zero-inflation model.

### Supplementary Tables

**Supplementary Table 1** (xlsx) - Sample metadata.

**Supplementary Table 2** (xlsx) - Trace gas oxidation rates for Dronning Maud Land *in situ* and *ex situ* incubations.

**Supplementary Table 3** (xlsx) - Microbial community metabolic marker genes (from short-read metagenomic data)

**Supplementary Table 4** (xlsx) - MAG quality, taxonomy, and metabolic genes.

**Supplementary Table 5** (xlsx) - Microbial community composition (from short-read metagenomic data)

**Supplementary Table 6** (xlsx) - Trace gas oxidation rates for Robinson Ridge, Dronning Maud Land, and Bunger Hills soil samples across 14 temperatures, plus Arrhenius and GTD-modelled values

**Supplementary Table 7** (xlsx) - Comparative analyses of modelled oxidation rates in the future under different emissions scenarios.

### Supplementary Figures

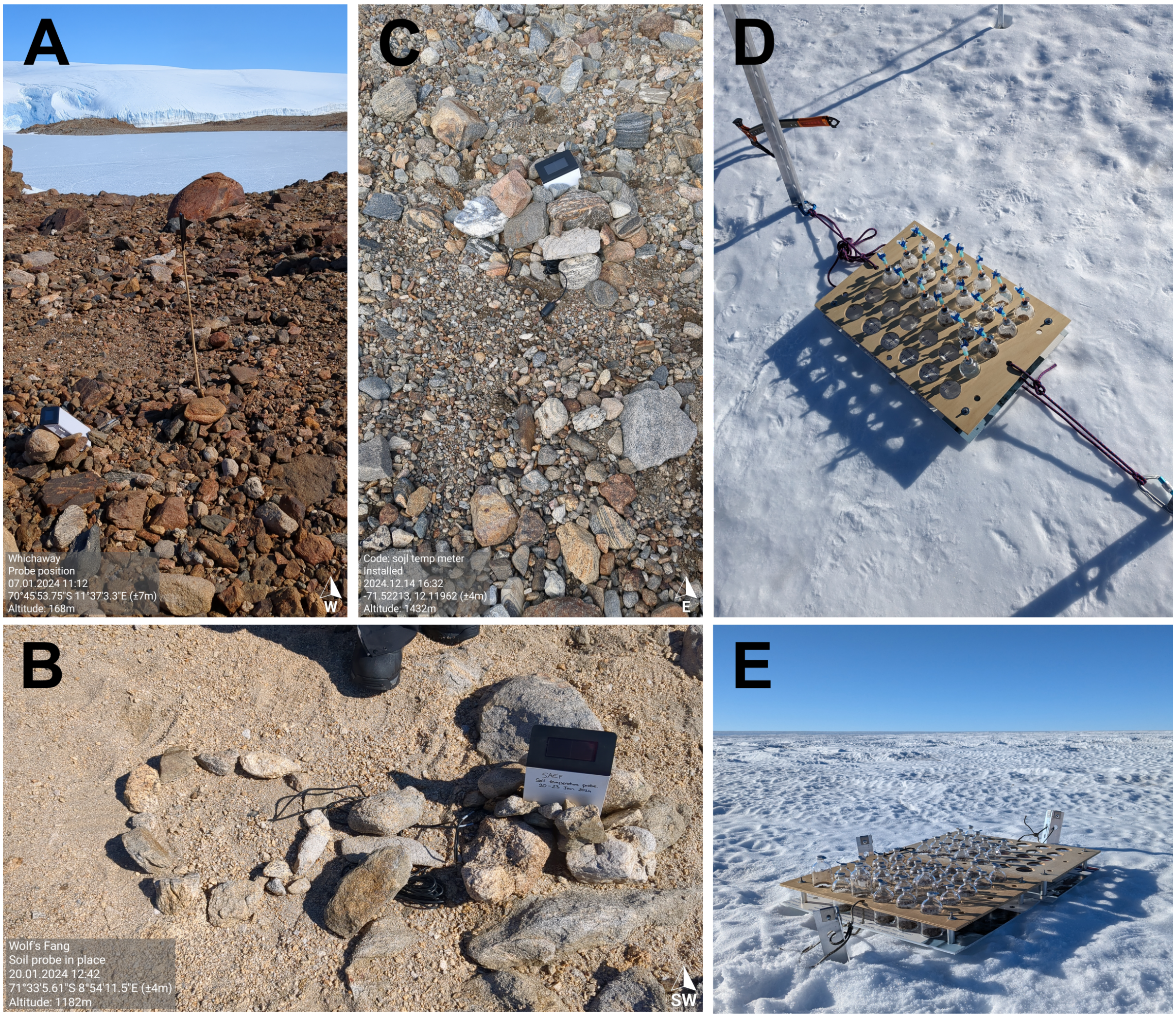


**Figure S1**. Locations where soil temperature was recorded at 10 cm depth in (A) Schirmacher Oasis, January 2024, (B) Henriksen Nunataks, January 2024, and (C) Petermann Ranges, December 2024. *In situ* incubations of Antarctic soil microcosms at field camps in Dronning Maud Land in (D) January 2024 and (E) December 2024.

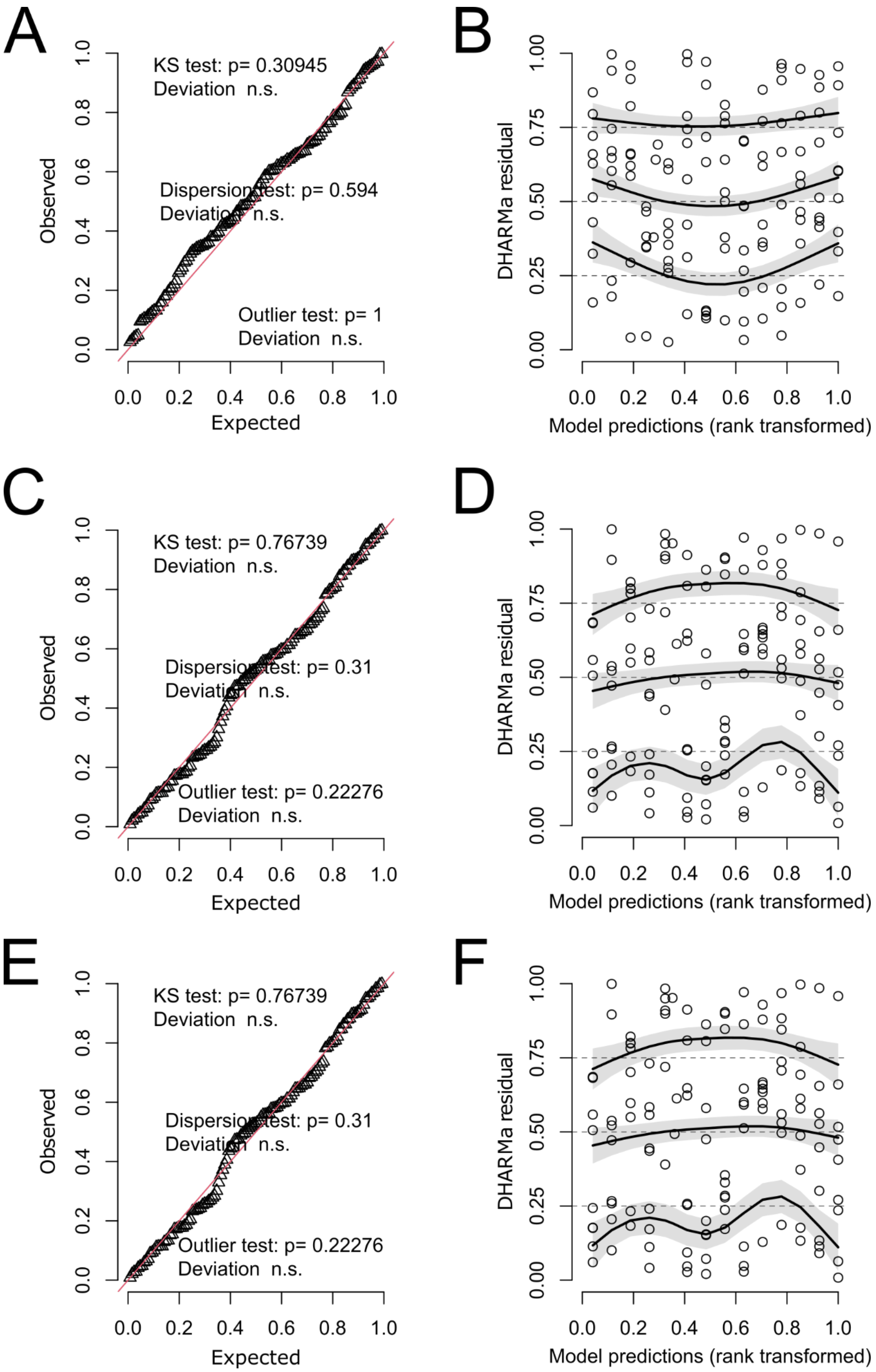


**Figure S2. Assessment of model fit and diagnostics.** QQ plot residuals for (**A**) $H_2$, (**C**) CO, and (**E**) $CH_4$. KS = DHARMa residual vs. predicted for (**B**) $H_2$, (**D**) CO, and (**E**) $CH_4$. No significant problems were detected in any of the residual vs. predicted plots.

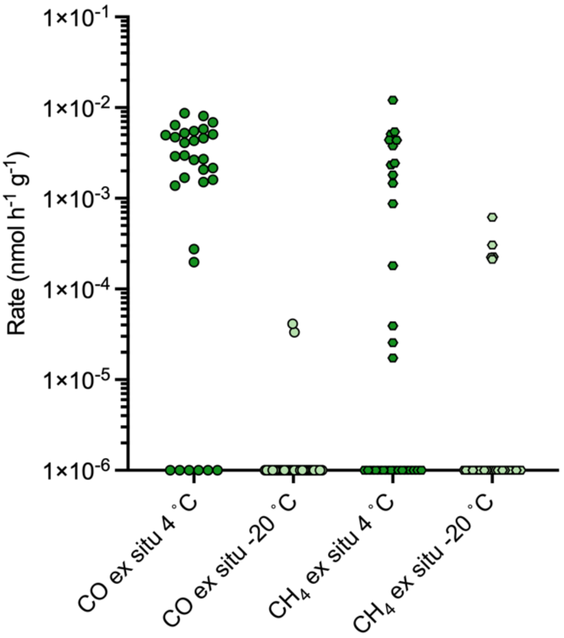


**Figure S3.** CO and $CH_4$ oxidation rates in *ex situ* microcosms of soils from Dronning Maud Land, Antarctica, incubated at 4 and -20˚C (*n* = 31). Samples with a no observable trace gas consumption (i.e. a rate of 0) are nominally shown at 1× $10^{-6}$ for representation on the log scale.

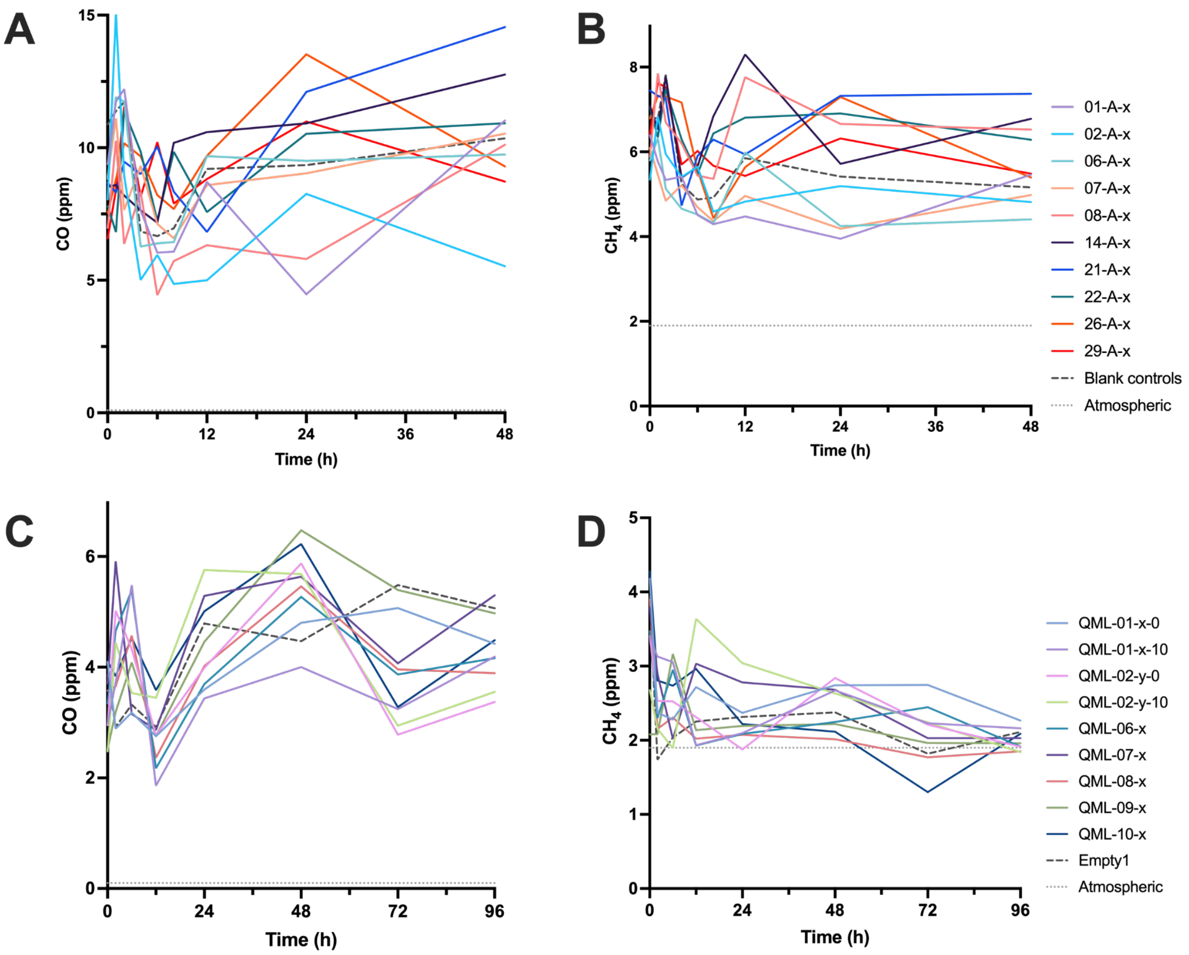


**Figure S4.** *In situ* incubations did not show conclusive CO or $CH_4$ consumption. (A) CO and (B) $CH_4$ measurements in 2023-24. (C) CO and (D) $CH_4$ measurements in 2024-25. All measured data points are triplicate averages. Error bars are not shown. Atmospheric concentration is also shown on each plot: 0.1 ppm for CO and 1.9 ppm for $CH_4$.

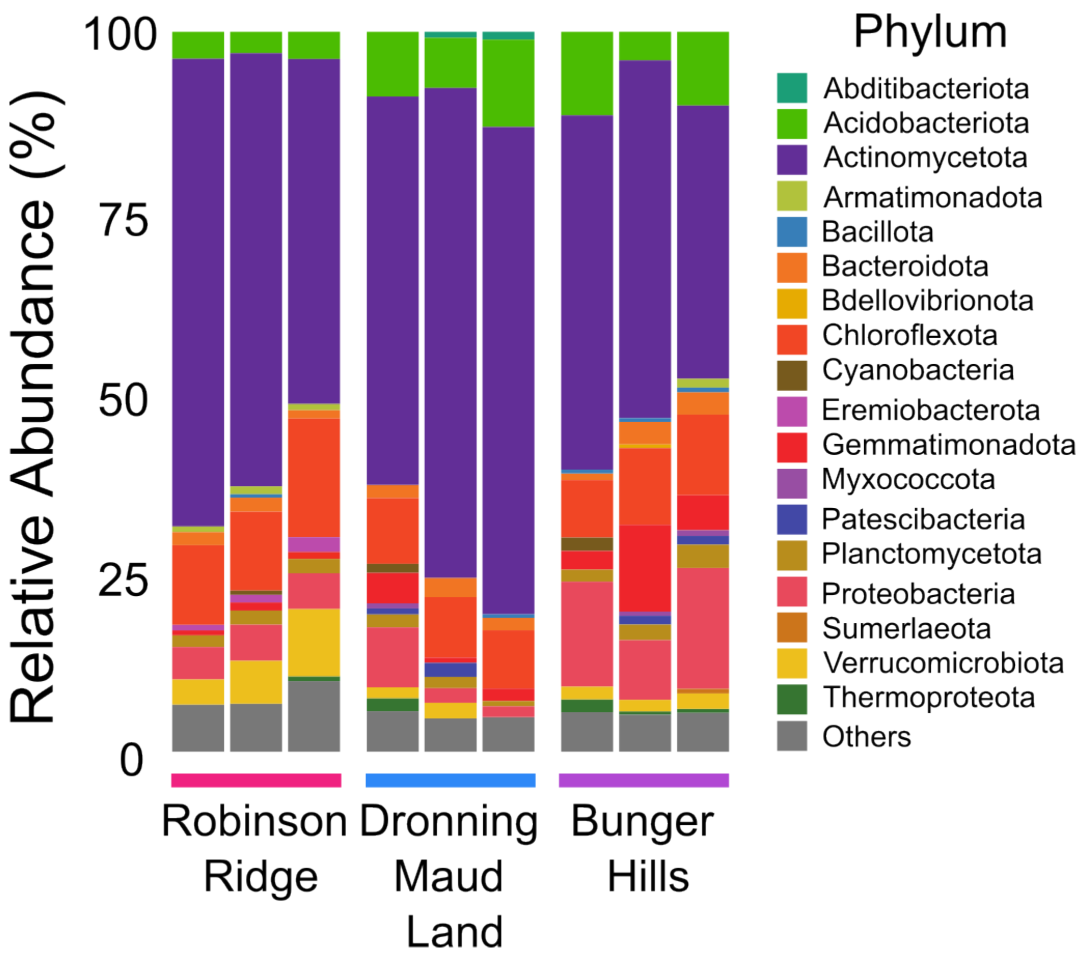


**Figure S5.** Taxonomic bar chart based on phyloFlash 16S rRNA gene taxonomic classification of the metagenomes. All phyla are bacterial except for Thermoproteota, which is archaeal. “Others” includes any bacterial and archaeal phyla with ≤100 counts.

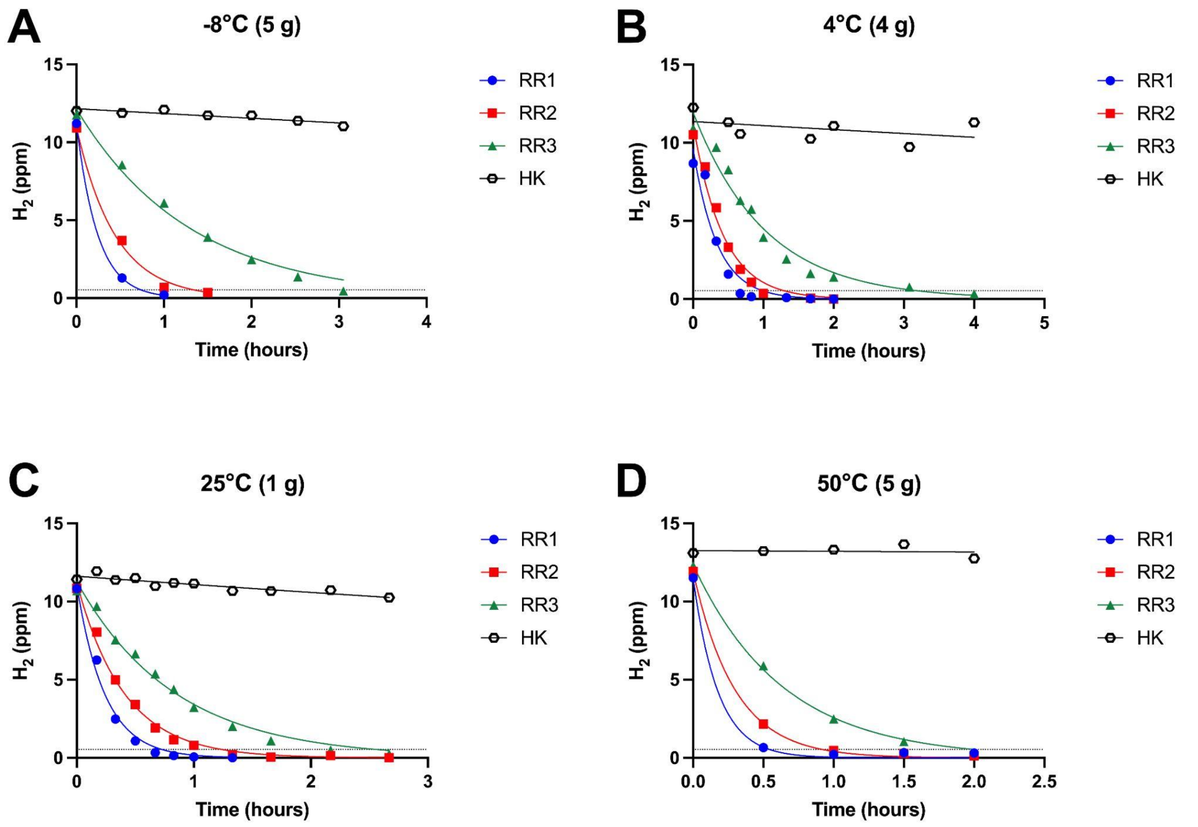


**Figure S6.** Select Robinson Ridge soils (RR1, RR2, RR3) and heat-killed controls incubated at (**A**) -8°C, (**B**) 4°C, (**C**) 25°C and (**D**) 50°C, showing $H_2$ oxidation to sub-atmospheric concentrations in less than 1 hour. The mass of soil used in each experiment is written in brackets in the graph title. Concentrations were fitted with non-linear regression models to derive rates. Dashed line indicates atmospheric $H_2$ concentration (0.53 ppm).

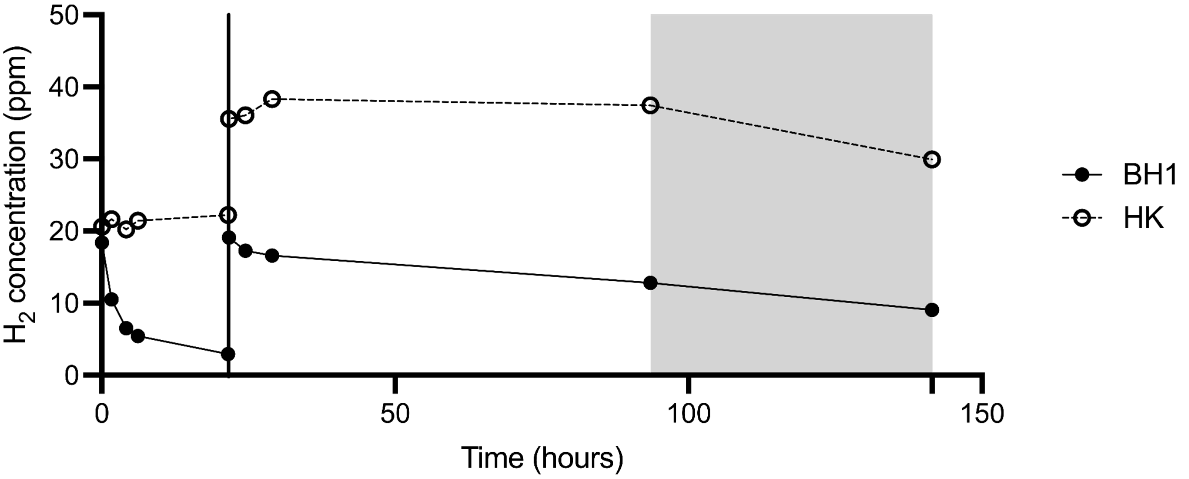


**Figure S7.** Bunger Hills (BH1) and heat-killed control (HK) soils at 75°C. Line at 24 h indicates injection of another ~20 ppm $H_2$. Shaded region represents the microcosm returning to room temperature (20°C).

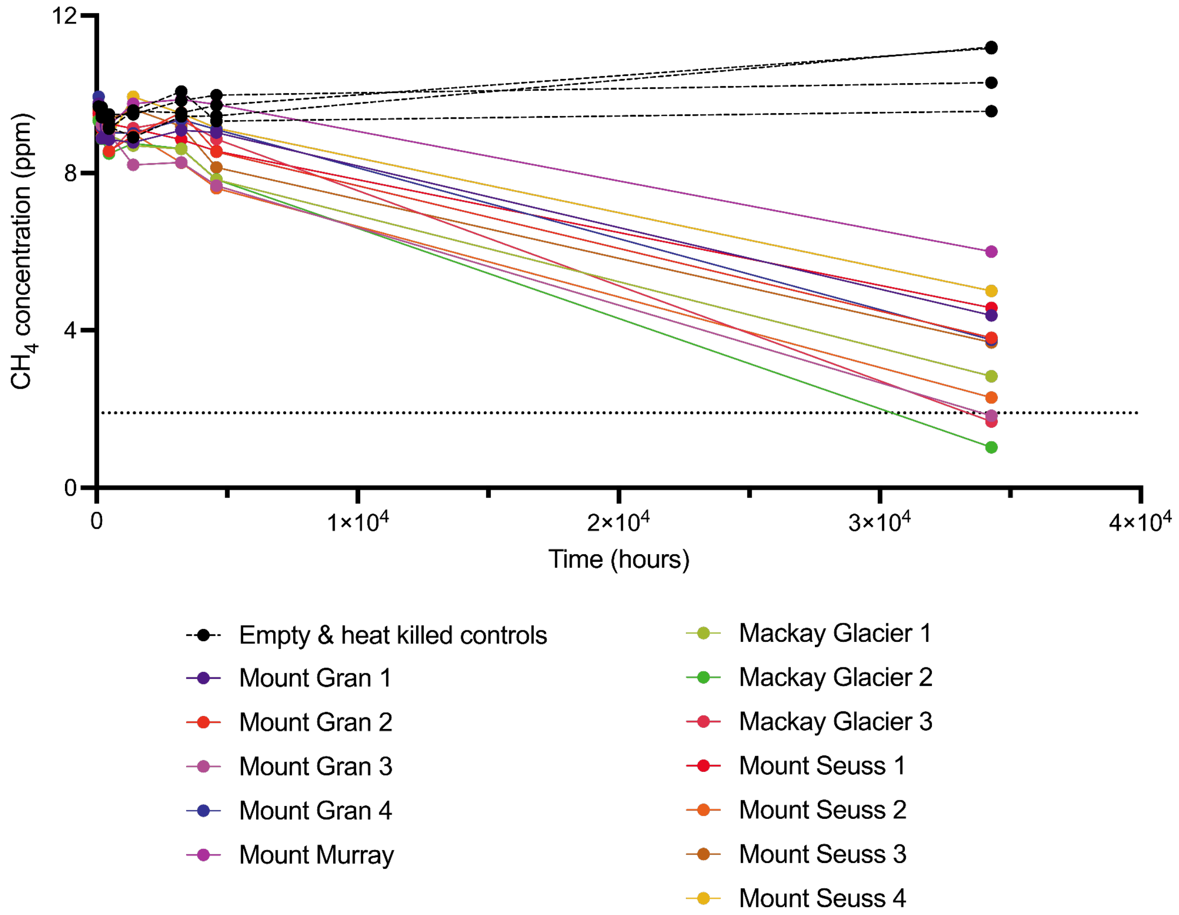


**Figure S8** Mackay Glacier soils and controls from Ortiz *et al.* (2021) were continuously incubated at -20°C. Data collected in the first ~6 months was published previously, while the final sampling occurred after ~4 years (34,273 h). Dotted line represents atmospheric $CH_4$ concentration (1.9 ppm).

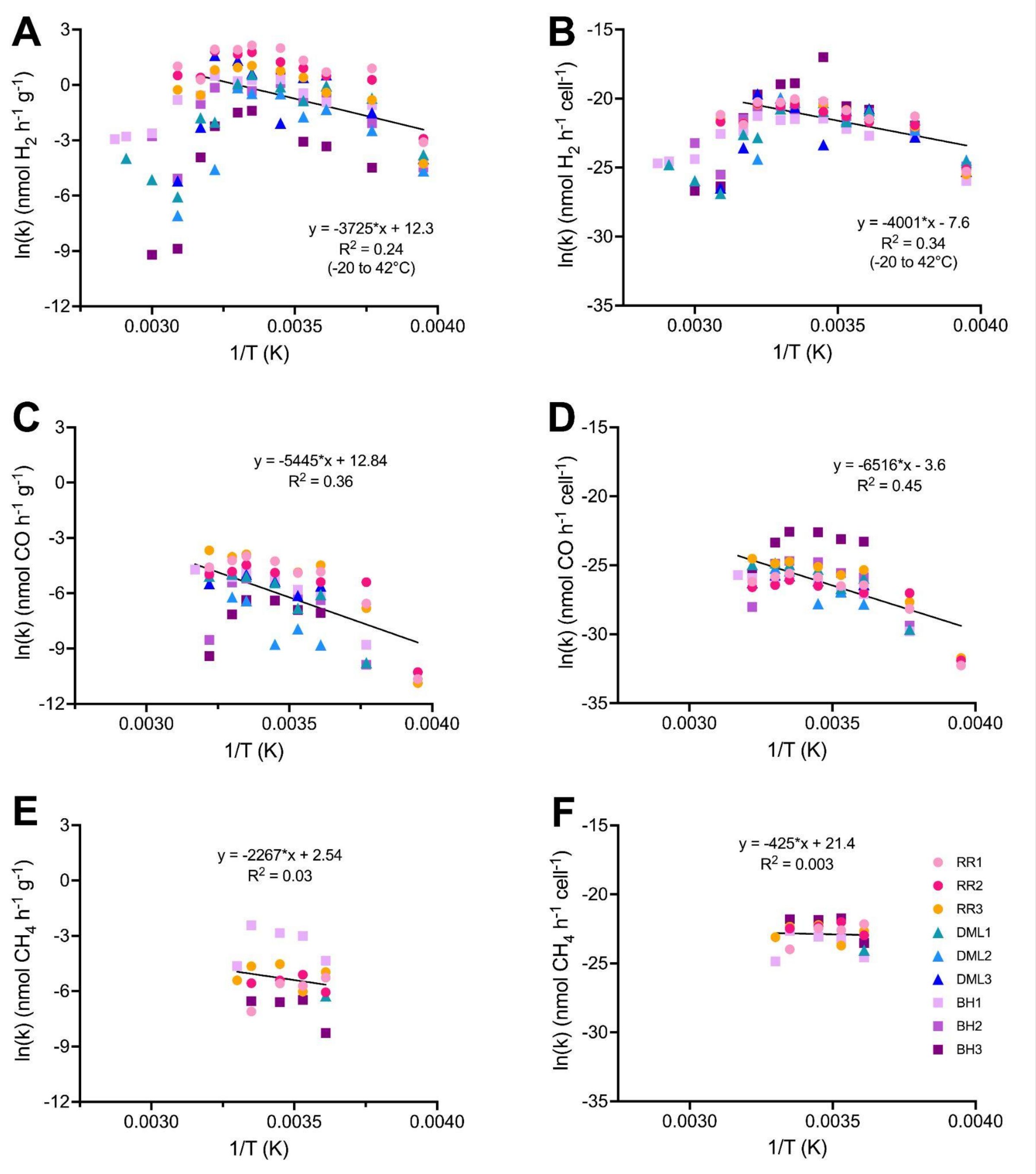


**Figure S9.** Arrhenius plots of both bulk oxidation rates (nmol $h^{-1}$ $g^{-1}$, **A,C,E**) and cell-specific oxidation rates (nmol $h^{-1}$ $cell^{-1}$, **B,D,F**). The Arrhenius equation was fitted to all soils from -20 to 42°C and is displayed in black. RR, Robinson Ridge; DML, Dronning Maud Land; BH, Bunger Hills.

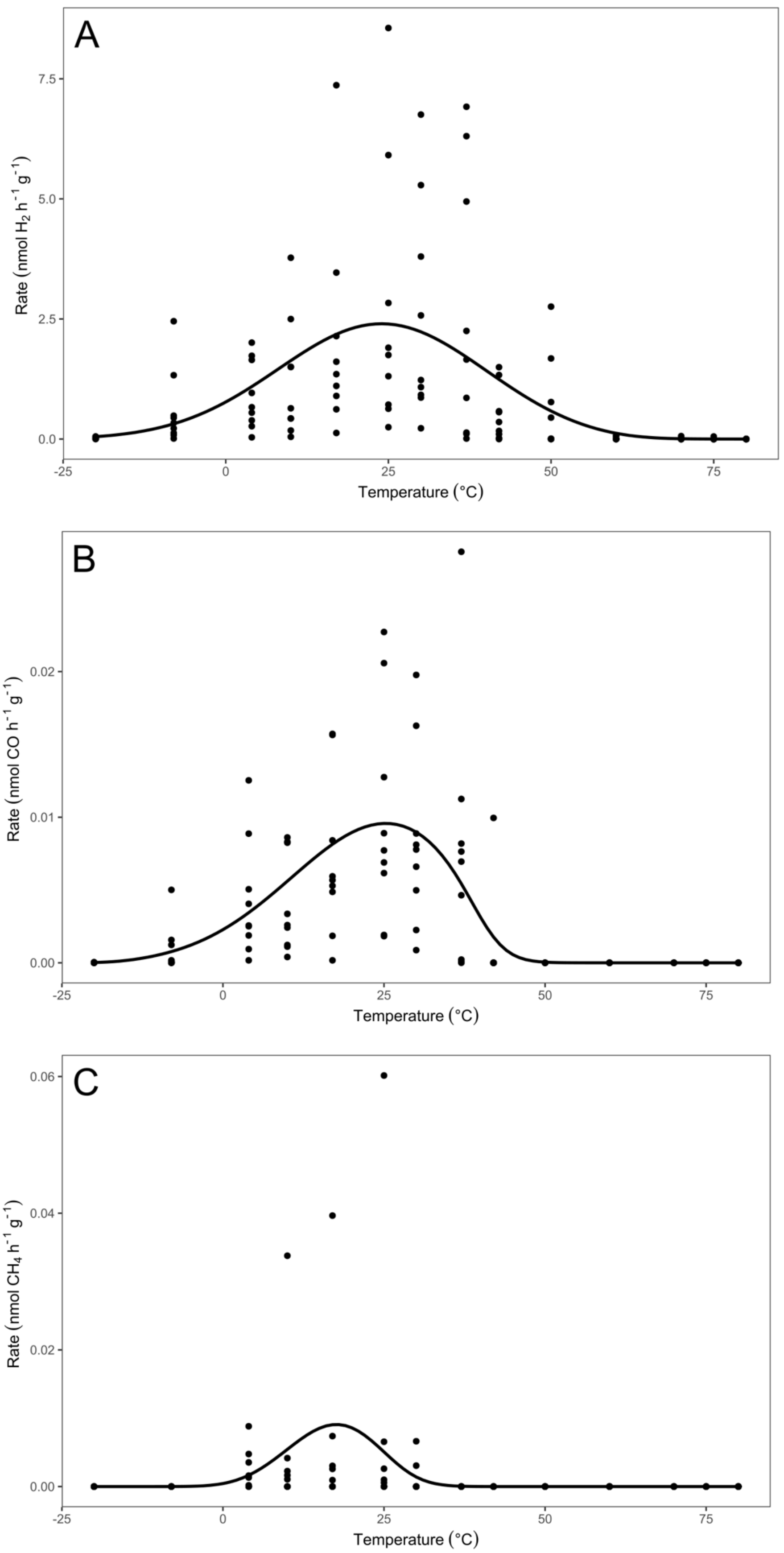


**Figure S10.** Generalised linear model for rates of (A) $H_2$, (B) CO and (C) $CH_4$ oxidation and temperature (°C).
